# Genome-wide ribonucleotide detection in *Archaea*

**DOI:** 10.1101/2025.03.17.643674

**Authors:** Yann Moalic, Maurane Reveil, Deepali L. Kundnani, Sathya Balachander, Taehwan Yang, Alli Gombolay, Farahnaz Ranjbarian, Raphael Brizard, Patrick Durand, Hannu Myllykallio, Mohamed Jebbar, Anders Hofer, Francesca Storici, Ghislaine Henneke

## Abstract

Genome integrity is constantly challenged by the incorporation of ribonucleotides (rNMPs) during DNA synthesis. Covalently linked single and several consecutive rNMPs occur in the genome of a number of organisms. They are mainly introduced by DNA polymerases during DNA replication and repair. In general, cells evolved ribonucleases H (RNases H) specialized in the removal of rNMPs from DNA to avoid any detrimental consequences on genome stability. Here, we describe the involvement of types 1 and/or 2 RNases H in processing embedded rNMPs in the genome of two archaeal species *Haloferax volcanii* and *Thermococcus barophilus*. Using combined approaches that include alkaline DNA fragmentation, high-throughput ribose-seq DNA sequencing and nucleotide pool quantification, the distribution, identity, level and sequence context of genomic rNMPs are reported and discussed regards to the intracellular balances of dNTPs and rNTPs. Our results confirm the predominant role of type 2 RNase H in the removal of genomic rNMPs. They also reveal rNMP-base compositions, densities, locations, and variations of surrounding bases at rNMP-embedment for each mutant. The cellular roles of the different RNases H in processing rNMPs in the genome of *Archaea* are discussed.

## INTRODUCTION

DNA replication is an essential process for all organisms. It consists of copying genetic information prior to cell division. Faithful replication of the genome requires that DNA polymerases bind and add the correct incoming dNTP to the growing DNA chain. The level of sugar discrimination and base selectivity depend on the DNA polymerase and the chemical identity of the template base (1,2). Because the level of RNA precursors (rNTPs) largely exceeds that of DNA precursors (dNTPs), DNA polymerases are constantly challenged with forming correct Watson-Crick base pairs (3–6). Despites having a “steric gate” for the exclusion of ribonucleotides, mistakes do occur (7). We recently showed that the archaeal family D DNA polymerase (PolD), also equipped with a steric gate (8), is able to pair or mispair single ribonucleotides opposite (deoxyribonucleoside monophosphate) dNMP-containing templates, and to pair or mispair single deoxyribonucleotides opposite (ribonucleoside monophosphate) rNMP-containing templates (4). Other studies also support the evidence of erroneous rNMP insertions in DNA which mainly occurs opposite a damaged DNA base in the template DNA by low-fidelity DNA polymerases (9–12). Therefore, rNMPs are commonly found in the genome of organisms. Moreover, up to a certain level, incorporated rNMPs in DNA can serve as positive signal for the cellular machinery. For example, they are involved in strand-discrimination during MisMatch Repair (MMR) in Eukaryotes (13,14) or in mating-type switching in *Saccharomyces pombe* (15). Nevertheless, excision of rNMPs from DNA is necessary to prevent excessive accumulation and deleterious effects, and cells have therefore evolved rNMP repair pathways. Single embedded rNMPs are primarily removed in an error-free manner by the evolutionary conserved Ribonucleotide Excision Repair (RER) pathway initiated by type 2 ribonuclease H (designated RNase HII in Prokaryotes and RNase H2 in Eukaryotes (3,16–18). In the absence of a functional RNase H2, Topoisomerase 1 (Top1) may operate as a backup for RER, which may be mutagenic or not (19–23). In Eukaryotes and *Bacteria*, other mechanisms ensure the removal of rNMPs like Base Excision Repair (BER) (24), MisMatch Repair (MMR) (25) and Nucleotide Excision Repair (NER) (26). Although the balance between incorporation and repair of rNMPs in DNA may vary across species, persistent rNMPs that exceed cellular tolerance, may lead to replication stress and genome instabilities (27–29). Indeed, high levels of genomic rNMPs due to defective RNase HII/2 are associated to disorders like Aicardi-Goutières (30,31) or Systemic Lupus Erythematosus (SLE) (32) in humans, tumorigenesis in mice (33–35) and mutagenesis in *Bacteria* (6,36). In addition to single genomic rNMPs, RNA:DNA hybrids (stretches of embedded RNA or R-loops) can also persist in DNA (for review, see reference (12)), and not only RNase HII/2 but also type 1 RNase H (designated RNase HI in Prokaryotes and RNase H1 in Eukaryotes) are involved in their removal (37,38). The specificity of RNase HI/1 differs from RNase HII/2 because it requires at least four consecutive rNMPs for cleavage (39–41), precluding any involvement in RER at least in Eukaryotes (16). Nevertheless, RNase HI from *Escherichia coli* (*E. coli*) is supposed to be involved in conjunction with RNase HII in RER (42). Another very recent work assigned a putative role of *E. coli* RNase HI in strand-specific removal of rNMPs, while the exact mechanism needs further clarification (43). Contrary to *rnhB* null cells (lacking RNase HII activity), which are asymptomatic, the absence of RNase HI *(ΔrnhA*) leads to clear phenotypes (*e.g*., smaller colonies, slower growth or unscheduled replication initiation) (37,44). Disruption of both RNase HI and RNase HII (*ΔrnhAB*) in *E. coli* revealed even greater effects (*e.g*., reduced viability or chromosome fragmentations), associated with accumulation of ribonucleotides in DNA and replication stress (37,45). On the other hand, RNase H1 is essential for mitochondrial DNA maintenance in mice (46), and RNase H1 deficient mutants result in human disorders like adult-onset (47,48).

In recent years, several sequencing-based technologies have been devoted to mapping ribonucleotides in DNA (49–55). This underlined the sequence-specific information about rNMP distribution in whole genomes, and the influence of rNTP/dNTP cellular ratio on rNMP incorporation. Notably, rNMP mapping has allowed defining replicative DNA polymerases usage genome wide in yeast (49,50,53), which revealed some rNMP incorporation patterns in different replicase-variant backgrounds (49,52), such as the non-preferential incorporation of rUMP. Besides, the use of RNR mutant cell lines demonstrated that dNTP pool imbalances drastically change rNMP incorporation profiles in yeast mitochondrial DNA (56). Evidence could also be made with the varying nucleotide pools, in which the high rATP/dATP ratio accounted for the vast majority of rAMP incorporation in genomic DNA of wild-type yeast cells (57). The sequence context was also studied and some preferred sites were depicted, suggesting potential base-stacking stabilization for the incoming rNTPs (57–59). Most reports in the literature referring to genome-wide profiling of rNMPs relate to yeast (49,50,52,53,57) and to a lesser extent to unicellular organisms (54,58).

Here, we decided to expand ribonucleotide mapping at nucleotide genome level in two archaeal models, *Haloferax volcanii* H53 and *Thermococcus barophilus* MP, using the ribose-seq technology (52) and Ribose-Map toolkit (60). While belonging to the same Euryarchaeota phylum, these two *Archaea* display some dissimilarities. *H. volcanii* (Hvo) is a halophile aerobe isolated from the Dead Sea (61) whereas *Thermococcus barophilus* (Tba) is a piezophile anaerobe hyperthermophile isolated from deep sea hydrothermal vents (62). Their optimal growth conditions are 45°C in 1.7-2.5 M NaCl and 85°C at 40 MPa, respectively. Their genome architecture is quite different (Table S1). *H. volcanii* H53 has one main circular chromosome (2.8 Mb), three plasmids pHV1 (85 kb), pHV3 (438 kb) and pHV4 (636 kb), while being cured of plasmid pHV2 (6 kb) (63). Compared to the GC-rich (65%) chromosome and plasmids of *H. volcanii*, the GC content is reduced in the chromosome (2 Mb) and pTBMP1 plasmid (54 kb) of *T. barophilus* MP (Table S1). Like for *H. volcanii* H53, *T. barophilus* MP laboratory strain is also cured of pTBMP1 (64).

Herein, we explored the role of RNases H in the occurrence of genomic rNMPs in *H. volcanii* strains lacking *rnhA* (Hvo_0732) or *rnhB* (Hvo_1978) or *rnhAB* (65) as well as in *T. barophilus* cells deleted for *rnhB* (TERMP_00671) (64). Moreover, the relevance of the flap endonuclease 1 (Fen1)(Hvo_2873) in rNMP removal was assessed for the triple knockout *ΔrnhAΔrnhBΔfen1 H. volcanii* cells (65). rNMPs in genomic DNA were revealed by alkaline hydrolysis and ribose-seq, a high-throughput DNA sequencing method which makes use of alkaline cleavage at rNMPs and ligation by *Arabidopsis thaliana* tRNA ligase (AtRNL) to exclusively capture 5’ rNMPs, while excluding abasic sites, nicks and Okazaki fragments (52). The identity and sequence context of rNMP incorporation were carefully inspected and discussed according to the intracellular balances of dNTPs and rNTPs.

## MATERIAL and METHODS

### Culture and harvesting of *Haloferax volcanii* and *Thermococcus barophilus* exponentially growing cells

*Haloferax volcanii* strains used in this study are the one used in Meslet-Cladiere (65). The strain H53 *(ΔpyrE2 ΔtrpA*) is referred as WT (Hvo WT). *H. volcanii* Δ*rnhA* (HvoΔ*rnhA*)*, H. volcanii ΔrnhB (*HvoΔ*rnhB*), *H. volcanii ΔrnhAΔrnhB* (HvoΔ*rnhA*Δ*rnhB*) and *H. volcanii ΔrnhAΔrnhBΔfen1* (HvoΔ*rnhA*Δ*rnhBΔfen1*) are respectively the strains CN10, CN7, CN11 and CN17. For accuracy and reminder, here is the list of the genes and their respective locus deletion on the chromosome: *pyrE2* gene (orotate phosphoribosyltransferase/ HVO_0333: 301,751-302,281) and *trpA* (tryptophan synthase subunit alpha/HVO_0789: 708,729-709,562) in all Hvo strains and, *rnhA* (HVO_0732: 654,302-654,895) and *rnhB,* (HVO_1978: 1,824,240-1,824,887) and *fen1* (HVO_2873: 2,710,411-2,711,391) in the mutant strains. Cells were grown at 42°C in YPC medium as previously described(63,65). 200 mL of cells in logarithmic growth phase at densities from 6.8 x 10^8^ to 27.4 x 10^8^ cells/mL were harvested by centrifugation, followed genomic or nucleotide extraction

*Thermococcus barophilus* MP (UBOCC-M-3107) and *T. barophilus ΔrnhB* (UBOCC-M-3303) cells were the two strains used in that study. For accuracy and reminder, here is the list of the genes and their respective locus deletion on the chromosome: For Tba Δ*rnhB*, the strain was constructed with the genetic tools (64), based on the strain deleted of the locus TERMP_00517: 432,339-432,986, *ΔrnhB* being at the locus TERMP_00671: 577,084-577,758.

Cells were grown in a gas-lift bioreactor under anaerobic conditions at 85°C in SME medium with a pH of 6.8 (64,66). Logarithmically growing cultures at a cell density of 2.8 x 10^8^ to 5.4 x 10^8^ cells/mL were harvested by centrifugation under strict anaerobia, followed by genomic extraction or nucleotide extraction.

### Nucleotide pool quantification

WT and RNase H mutant *H. volcanii* cells were collected by centrifugation for 5 min at 6000 g at 4°C. Cell pellets were washed with 5 ml ice-cold NaCl solution (NaCl 144 g/L) before centrifugation at 6000 g for 2 min at 4°C. The supernatants were discarded and cell pellets submitted to nucleotide extraction. WT and RNase H mutant *T. barophilus* cells were harvested by centrifugation at 15000 g for 5 min at 4°C. Each cell pellet was immersed in 100 µl ice-cold NaCl solution (NaCl 14 g/L, 2% glucose) under anaerobic conditions before nucleotide extraction. The immersion step was included for the *T. barophilus* cells because it helped to increase the final yield of the nucleotides. The nucleotide extraction protocol was adapted from Hofer et al., 1998(67). For *H. volcanii* strains, cell pellets were resuspended in 500 µL of ice-cold 12% (w/v) trichloroacetic acid (TCA) and 15 mM MgCl_2_ solution. For *T. barophilus* strains, 100 µl ice-cold 24% (w/v) TCA and 30 mM MgCl_2_ solution was added to each tube (containing 100 µl NaCl-glucose solution) and the content was mixed under anaerobic conditions by pipetting the cells up and down. The cells were flash-frozen in liquid nitrogen and incubated for 15 min with stirring at 4°C. This flash-freezing step was performed one time for *H. volcanii* strains and three times for *T. barophilus* strains. In the case of *T. barophilus*, the content of three tubes was pooled resulting in a sample volume of 600 µL.

The supernatant of each sample was collected by centrifugation for 1 min at 18 000 g at 4°C and extracted in 750 µL of Freon-Trioctylamine solution (75.2% v/v Triochlorotrifluorethane and 21.8% v/v trioctylamine). After centrifugation, the aqueous phase (upper phase) containing the nucleotides was collected and subjected to a second extraction with 500 µL of Freon-Trioctylamine solution. The upper phase was saved and the pH was confirmed to be above 5 by spotting 0.2 µl on a pH paper. The resulting solution was finally purified by solid phase extraction as described before prior to HPLC analysis (68).

Nucleotide analysis was performed by the Fast HPLC protocol in which ADP, and all eight NTPs and dNTPs are separated in a single run on a reverse phase column in the presence of tetrabutylammonium ions as ion pairing agent (68). The quantification of the nucleotides was then performed by comparing the peak areas for abundant nucleotides (ADP and NTPs) and peak heights for dNTPs to a nucleotide standard.

### Genomic DNA extraction

Logarithmically growing cells were harvested by centrifugation for 1 h, 8000 g at 4°C. Cell pellets (∼0.5 μg) were suspended in 1.6 mL TE [10 mM Tris–HCl (pH 8), 1 mM Na_2_EDTA]. To ensure cell lysis, 100 μL proteinase K (20 mg/mL), 200 μL Sarkosyl (10%) and 200 μL SDS (10%) were added followed by subsequent incubation at 37 °C for 1.5 h. Isolation of total DNAs was accomplished by adding an equal volume of buffered (pH 8.0) Phenol/chloroform/isoamylalcohol (25:24:1). The DNA solution was gently mixed and the aqueous phases were collected by centrifugation at 10,000g for 10 min at 4 °C (done three times). RNA degradation was performed with 0.05 µg/µL RNase A in TE with high salt (0.5 M NaCL) at 37°C for 1 h. DNAs were purified with an equal volume of phenol/chloroform/isoamylalcohol and centrifuged. The upper phase was extracted with an equal volume of pure chloroform (25:24:1) and centrifuged. DNA precipitation was obtained by mixing the final aqueous phase with a 0.7 volume of 100% isopropanol followed by incubation for 1 h at room temperature. After a 30-min centrifugation at 15,000g at 4 °C, the DNA pellets were washed once with 0.5 mL of 70% ice-cold ethanol. Finally, the DNA pellets were air-dried for 45 min before suspension in nuclease-free water.

### Detection of rNMPs in genomic DNA

Detection of genomic rNMPs in the genomes of wild-type and RNase H null archaeal cells were achieved as previously described (4). Given that alkali hydrolysis may also reveal local nicks, breaks and gaps, a control experiment was also performed with the genomic DNA treated with NaCl instead of NaOH. This treatment was followed by formamide denaturation before agarose gel electrophoresis. This formamide DNA denaturing method allows for the separation of ssDNA without hydrolysing rNMP phosphodiester bonds (69). DNA separated under non-denaturing conditions allow the identification of the intact genomic DNA. Briefly, ∼10 µg of DNAs were heated for 2 h at 55°C with either 0.3 M NaOH or 0.3 M NaCl. Samples were ethanol precipitated and dissolved in either 90% formamide, 20 mM EDTA (denaturing conditions) or neutral buffer (TE, 30% glycerol) (neutral conditions), followed by separation on 1% agarose gels and by staining with ethidium bromide. Densitometry traces were obtained with Quantity One software 4.6.6 (Bio-Rad). Each lane profile was extracted from the same gel and the background subtracted. The quantification of DNA fragmentation pattern was calculated from densitometry traces as follows: densitometry intensity distribution was divided by the fragment length distribution to quantify the proportion of molecules of a particular fragment size.

### Ribose-seq library preparation

The ribose-seq libraries were prepared as previously published with the following changes (52,57,58). DNA was fragmented by two sets of restriction enzymes (RE) with or without dsDNA Fragmentase (New England Biolabs): RE1 (*DraI*, *EcoRV*, and *SspI*) and RE2 (*MspA1I*, *HpyCH4V*, and dsDNA Fragmentase). Each cocktail of RE was incubated with DNA at 37°C overnight and generated DNA fragments up to 3,000 bp with an average size of 1,500 b. In the case of dsDNA Fragmentase, it was incubated with DNA at 37°C for 30 minutes, generating DNA fragments up to 1,000 bp. A total of two rounds of ribose-seq libraries were prepared (R1 and R2). In the PCR steps, R1 libraries (FS98 ∼ FS102) were amplified for 15 cycles and 10 cycles in the first and second PCR rounds, respectively. R2 libraries (FS217 ∼ FS222) were amplified for 10 cycles in both PCR rounds. The constructed libraries were sequenced on the Illumina NextSeq platform.

### Processing and alignment of sequencing reads

The ribose-map toolkit (60) was used to generate the BED files containing the genomic coordinates of the embedded rNMPs (Modules “alignment” and “coordinate”). Following the same process as previous studies (57,70), the background noise of restriction enzyme reads as well as the reads containing dAMP, which correspond to the dA-tailing step of the ribose-seq protocol, were removed (71).

### Nucleotide sequence context of embedded rNMPs

For each rNMP incorporation site, the sequence surrounding of 100 nucleotides upstream and downstream was extracted from the reference genome of each species. Then, the frequencies of nucleotides were calculated at each position. This was calculated for each targeted nucleotide separately and with all combined nucleotides. The results presented are normalized according to the reference proportion of each nucleotide in their respective genome (Chromosome and plasmids for *H. volcanii and T. barophilus*) as in previous studies of ribose-seq (52,57,58).

### Heatmaps

The mononucleotide and dinucleotide (RN and NR) heatmaps were generated as in Balachander *et al*. (57). In brief, the number of each type of rNMP (R_N_: R_A_, R_C_, R_G_, or R_U_) was counted and divided by the corresponding background of deoxyribomononucleotide of the reference genome to get frequencies that are then normalized after its division by the sum of the all the four frequencies. The normalized frequency was calculated using the RibosePreferenceAnalysis package(70).

## RESULTS

### Genome-wide accumulation of rNMP sites in RNase HI and RNase HII mutants

The DNA of the different mutant strains was used as templates for the robust genome-wide mapping technology ribose-seq (52). Two rounds of ribose-seq (R1 and R2) were performed with two sets of restriction enzymes (RE1 and RE2) in order to assess the GC percent influence on the methodology and whether fragmentation bias might occur. As observed in Table 1, the GC content of the chromosomes is reverted between each model archaeal strain (66.7% for Hvo and 42% for Tba). While the first RE set (RE1) assigned to ribose-seq library preparation was biased towards AT-rich regions, the second set (RE2) brought a better balance between AT- or GC-regions, ensuring a higher detection of rNMPs for all samples (Table 1 and Table S2). These results were supported by *in silico* restriction analysis of chromosomes in *H. volcanii* and *T. barophilus* strains showing a higher density and coverage of target DNA at RE2 enzyme-recognition sites (dsDNA Fragmentase not included) (Figure S1). This increase is even more pronounced for the chromosome of *H. volcanii* than the one of *T. barophilus* (Figure S1).

**Table 1.**
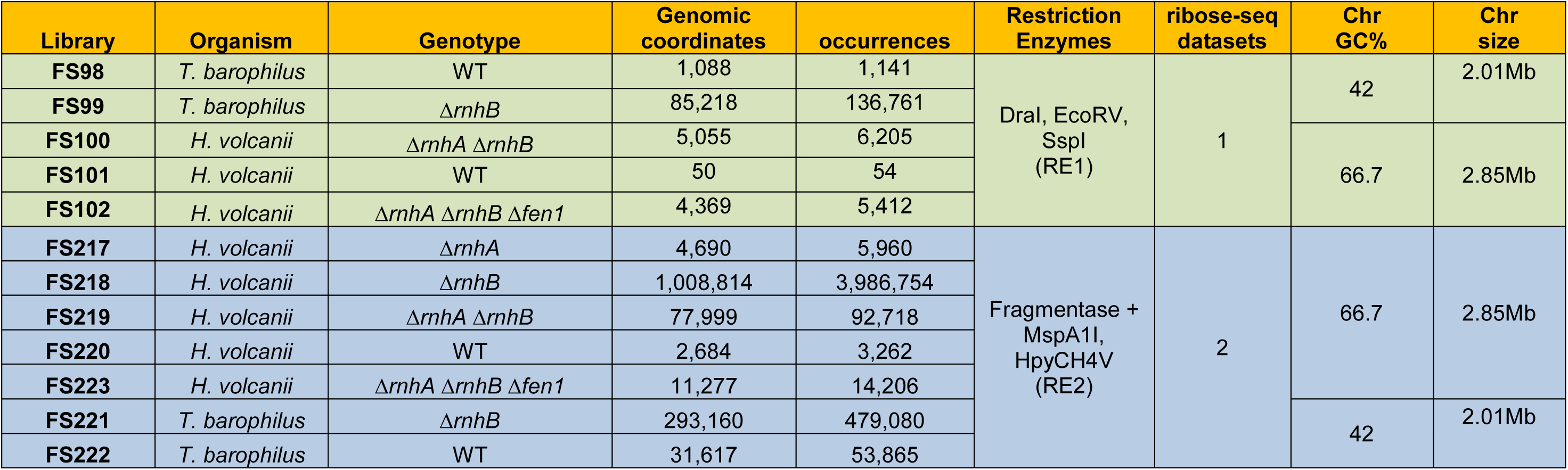
ribose-seq datasets generated for this study. Genomic coordinates are the number of different locations of rNMPs detected on the chromosome while occurrences are the overall rNMPs detected. The higher values of occurrences mean that several rNMPs are detected for the same genomic coordinates. RE1 = Restriction Enzymes set 1. Cut Sites: DraI = TTT/AAA, EcoRV = GAT/ATC, SspI= AAT/ATT. RE2 = Restriction Enzymes set 2. Cut Sites: MspA1I = CMG/CKG, HpyCH4V = TG/CA. Fragmentase is a non-specific endonuclease used in NGS library constructions (84).

We then examine the rNMP content into the genome of all archaeal strains by alkali treatment followed by agarose gel electrophoresis. Alkali causes strand cleavage at embedded rNMPs in DNA, resulting in a smear of products that runs down toward the bottom of the gel as well as a loss of the intact genomic DNA band (see legend in Figure 1 and methods for more details). As expected, irrespective of the RE cocktails used, the level of genomic rNMPs increased in strains lacking RNases H (Figure 1). In particular, DNA mobility of Hvo *ΔrnhA*, Hvo *ΔrnhAΔrnhB* and Hvo *ΔrnhB* gradually increased after alkaline treatment regards to Hvo WT (Figure 1A, compare lanes 5, 8 and 11 with 2), giving a clear peak shift of densitometry trace for Hvo *ΔrnhB* (Figure 1A, densitometric green line). Interestingly, compared to Hvo *ΔrnhB*, the loss of RNase HI in the double mutant *ΔrnhAΔrnhB* slightly reduced retention of genomic DNA (Figure 1A, compare lanes 8 and 11). Moreover, total genomic rNMPs increased in the triple mutant Hvo *ΔrnhAΔrnhBΔfen1* compared to Hvo WT (Figure 1B, compare lanes 2 and 8). However, in contrast with Hvo *ΔrnhAΔrnhB*, lower embedded rNMPs were detectable in Hvo *ΔrnhAΔrnhBΔfen1,* as shown by the reduction in alkali-sensitivity of genomic DNA (Figure 1B, compare densitometric orange and blue lines). The calculated rNMP frequencies also supported the highest fragmentation patterns observed for Hvo *ΔrnhB* (∼1 rNMP every 900 bp±50, Figure 1D) compared with Hvo *ΔrnhAΔrnhB* (∼1 rNMP every 1150 bp±71 or 1240 bp±14, Figure 1D, R1 and R2, respectively), Hvo *ΔrnhA* strains (∼1 rNMP every 1350 bp±71, Figure 1D), and Hvo *ΔrnhAΔrnhBΔfen1* (∼1 rNMP every 1530 bp±71, Figure 1D). As expected, the genome of Tba *ΔrnhB* cells displayed more alkali-sensitive sites than Tba WT cells (Figure 1C, lane 2 compared to lane 5), giving a substantial shift of densitometry trace (Figure 1C, densitometric green line compared with the red). The estimated incorporation frequency in genomic DNA was ∼1 rNMP every 830 bp in Tba *ΔrnhB* compared to ∼1 rNMP every 1460 bp in Tba WT (Figure 1D).

**Figure 1.**
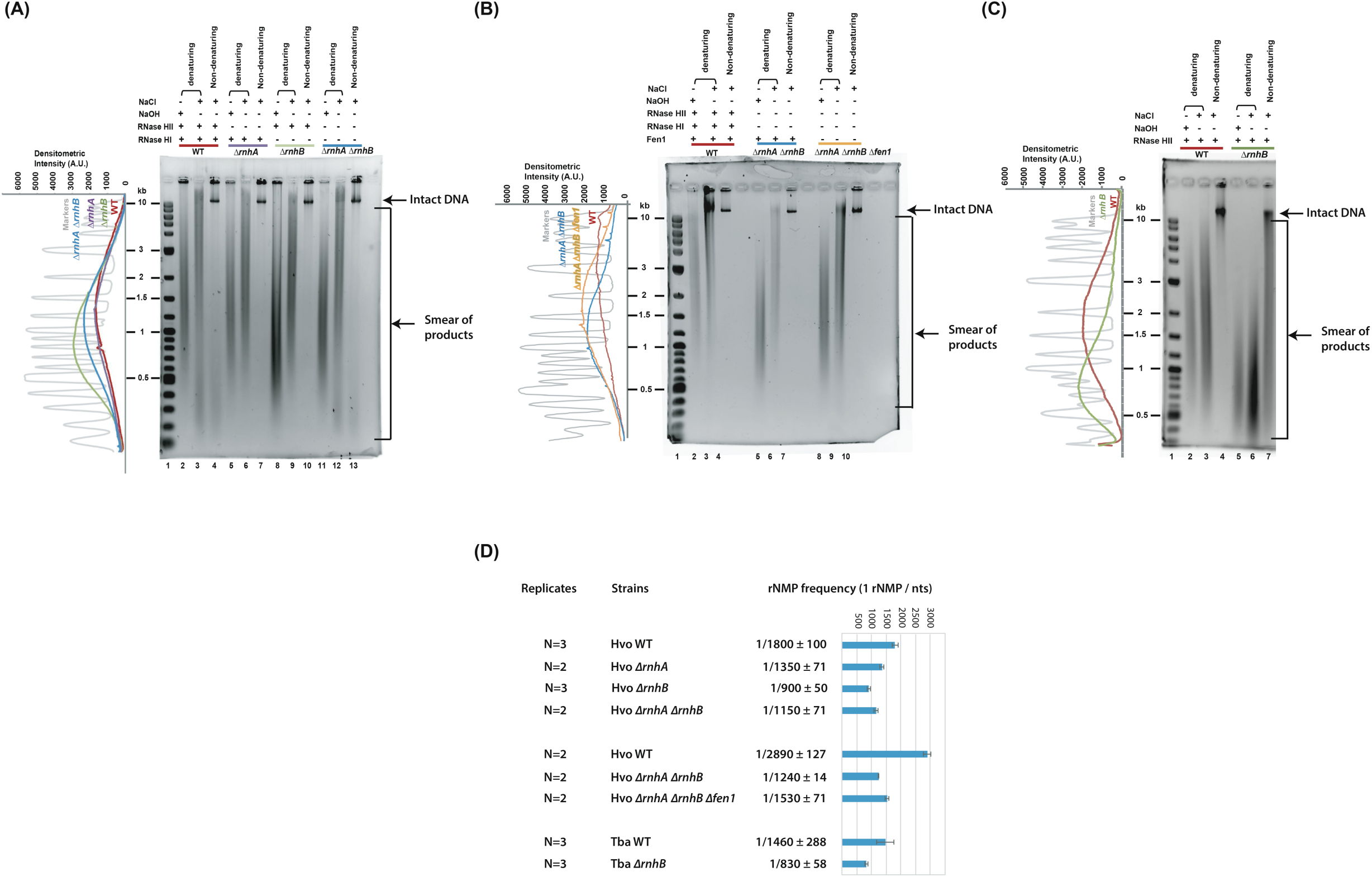
Alkaline hydrolysis of the genomic DNA from exponentially growing archaeal cells. Separation by agarose gel electrophoresis under neutral conditions (non-denaturing) and after denaturation with formamide (denaturing) followed by densitometry analysis of NaOH-treated DNA. Size of the DNA markers is indicated. **(A)** Alkali sensitive sites in genomic DNA of wild-type (WT), Δ*rnhA*, Δ*rnhB* and Δ*rnhA*Δ*rnhB* strains of *H. volcanii*. Densitometry intensity is plotted on the left for the selected lanes 2, 5, 8, and 11. **(B)** Alkali sensitive sites in genomic DNA of wild-type (WT), Δ*rnhA*Δ*rnhB* and *ΔrnhAΔrnhBΔfen1* strains of *H. volcanii*. Densitometry intensity of lanes 2, 5 and 8. **(C)** Alkali sensitive sites in genomic DNA of wild-type (WT) and Δ*rnhB* strains of *T. barophilus*. Densitometry analysis of lanes 2 and 5. **(D)** Averaged frequency of genome-embedded rNMPs of NaOH-treated samples from (A), (B), and (C).

On the other hand, the occurrence of rNMPs in RNase H mutants relative to RE1 and RE2 cocktails (Table 1 and Table S2) gave consistent results with the alkali-fragmentation of total genomic DNA in all strains (Figure 1A-C). Genome-wide density of rNMPs in the chromosome and genetic elements (pHV1, pHV3 and pHV4) of *H. volcanii* strikingly increased with the RE2 sequencing datasets (Figure 2A and Figure S2, compare Hvo WT-R1, *ΔrnhAΔrnhB*-R1 and *ΔrnhAΔrnhBΔfen1*-R1 to Hvo WT-R2, *ΔrnhA*-R2*, ΔrnhB*-R2, *ΔrnhAΔrnhB*-R2 and *ΔrnhAΔrnhBΔfen1*-R2). Moreover, the rNMP occurrence gradually decreased upon successive deletions of RNase HI and Fen1 in the *ΔrnhB* mutant (Figure 2A and Figure S2, R2 datasets), which is consistent with the DNA mobility patterns observed in Figure 1A-B. The highest enrichment of rNMPs across the chromosome and plasmids was always identified in *ΔrnhB* (Figure 2A and Figure S2), agreeing with the increased alkali-fragmentation of total genomic DNA (Figure 1A-B). On the other hand, a clear correlation between the size of DNAs and the prevalence of rNMPs was found, in which the number of rNMP genomic coordinates in the chromosome surpassed that of plasmids (Figure 2A compare with Figure S2). Genome-wide mapping of rNMPs in *T. barophilus* revealed an elevated level of rNMPs, and the loss of RNase HII clearly favoured their retention throughout the chromosome (Figure 2B and Table 1, compared Tba WT and Tba *ΔrnhB* in R1 or R2 datasets). This result is clearly consistent with the increased DNA mobility of Tba *ΔrnhB* after alkali treatment compared to Tba WT (Figure 1C). Comparison of the genome-wide accumulation of rNMPs across wild-type archaeal genomes indicated a higher enrichment for *T. barophilus* cells (Figure 2A-B, compare Tba WT-R2 and Hvo WT-R2 datasets; Table 1, compare rNMP occurrences Tba WT and Hvo WT for R2 datasets). However, depletion of RNase HII dramatically exacerbated the occurrence of rNMPs throughout the chromosome of *H. volcanii* compared with *T. barophilus* cells (Figure 2A-B, compare Tba *ΔrnhB*-R2 and Hvo *ΔrnhB*-R2 datasets; Table 1, compare rNMP occurrences Tba *ΔrnhB* and Hvo *ΔrnhB* for R2 datasets).

**Figure 2.**
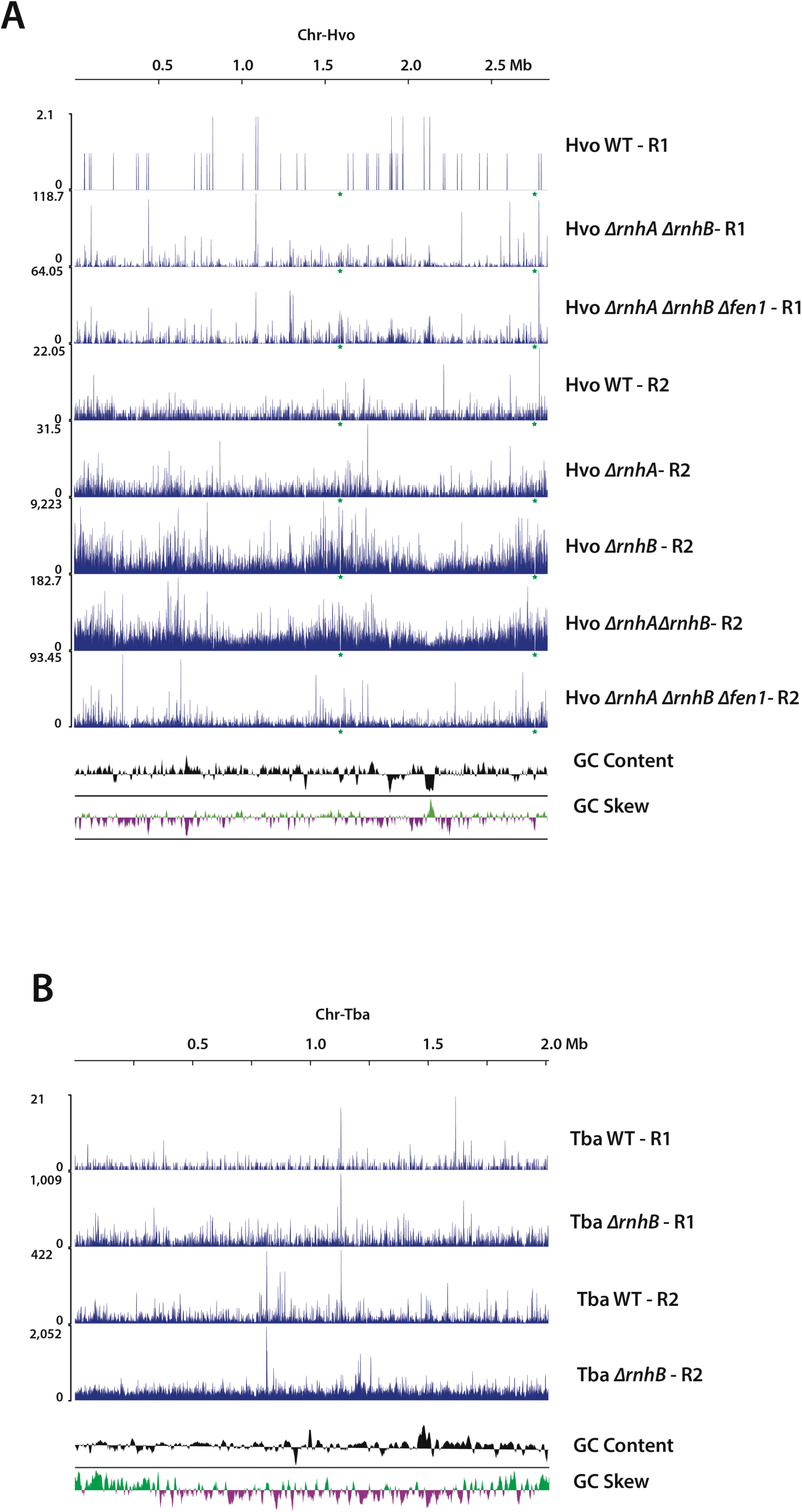
Chromosome-wide profiles of ribonucleotide incorporation. Locations and occurrences of rNMPs extracted from two rounds of ribose-seq (R1 and R2) on the chromosomes of *H. volcanii* (Chr-Hvo) **(A)** and *T. barophilus* (Chr-Tba) **(B)**. Each profile represents a mutant or a wild-type (WT) strain of these organisms. Occurrence of rNMPs is plotted in every 1 kb-sized chromosome sequences. The green stars give the positions of the coding locus of 16S and 5S rRNA that show no incorporation of rNMP in any of Hvo strains. Panels **(C)** and **(D)** illustrate the strong difference between the same coding locus of 16S and 5S rRNA that harbour no rNMP incorporation for Hvo Δ*rnhB* (C) and a strong level of rNMP incorporation for Tba Δ*rnhB* (D). Panels C and D are extracted from Proksee (https://proksee.ca/)(83).

Regards to the forward and reverse DNA strands, no strong differential rNMP incorporation was identified between R1 and R2 datasets across the chromosomes of *H. volcanii* and *T. barophilus* when a sufficient number of rNMPs was detected (Table S3 and Table S4, respectively). Nevertheless, a preferential rNMP segregation to the reverse strand in the chromosome of all *H. volcanii* strains was slightly noticeable, whereas strand-bias preference toward the forward strand was found in most plasmids (Table S3 and Table S4). For *T. barophilus* cells, the incorporation values showed a higher accumulation in the forward strand of Tba WT-R2 that shifted toward the reverse strand of Tba *ΔrnhB*-R2 (Table S3). Regards to R1 datasets, a strong bias toward the forward strand of the two genotypes was observed (Table S4).

### Genome-wide fine mapping of rNMP sites in RNase HI and RNase HII mutants

Analysis of the global distribution of rNMP sites across the genomes (Chromosomes and plasmids) revealed underrepresented (rNMP coldspots) or overrepresented (rNMP hotspots) regions (Figure 2 and Figure S2, see RE2 datasets). However, no relationship between hotspots or coldspots and GC content or GC skew profiles was found (Figure 2 and Figure S2, see roller coaster patterns). Then, we investigated chromosome-based features at rNMP hot/cold spots in *H. volcanii* and *T. barophilus* strains.

The results showed characteristic coldspots in the chromosome of *H. volcanii* cells, which were clearly visible with R2 datasets (Figure 2A, see green stars). Our fine-scale analysis ascribed mainly coldspots spanning the chromosomal coordinates 1,597,828 to 1,603,136 (locus Hvo_RS13015-25) and the coordinates 2,766,690 to 2,771,988 (locus Hvo_RS18910-20), which encode the 16S and 23S ribosomal RNA genes. Conversely, rNMP hotspots at the 16S ribosomal RNA (16S rRNA) gene (coordinates 812,419 to 817,064 or locus TERMP_RS04670-80) have been identified in the chromosome of Tba *ΔrnhB* strain. Harbouring 2253 rNMPs, this hotspot region spans 3 kb and contains flanking sequences of densely and sparsely clustered rNMPs at starting and ending positions of the 16S rRNA gene, respectively. The same hotspot region and distribution of rNMPs across the 16S rRNA gene was also observed in the chromosome of WT Tba.

Examining the strand specificity of rNMP distribution across the chromosomes with a Gaussian filter for R2 datasets allowed the identification of particular patterns in *H. volcanii* strains lacking RNase HII (Figure 3). Indeed, the red (forward strand) and blue (reverse strand) curves intercepted at some specific genomic coordinates in Hvo *ΔrnhB* (Figure 3A), overlapping with the previously described chromosomal replication origins *oriC1*, *oriC2* and *oriC3* (0 kb, 571 kb and 1593 kb, respectively)(72). At these origins (vertical dashed lines, Figure 3A), the distribution of rNMP was strand-specific in Hvo *ΔrnhB*. On the forward strand (red curve, Figure 3A), rNMP incorporation was favoured just upstream each *oriC*, but then was immediately reversed downstream. In this strain, the increased density of rNMP incorporation at the *oriC* was coincident with origin firing (72). Like Hvo *ΔrnhB*, the pattern of rNMP incorporation for Hvo *ΔrnhAΔrnhB* and Hvo *ΔrnhAΔrnhBΔFen1* strains (R2) was maintained with curves crossing at the three main chromosomal origins, but with lower amplitude (Figure 3A). However, these *oriC*-intersecting profiles were not detectable with Hvo WT and Hvo *ΔrnhA* (Figure 3A). Furthermore, no curve crossing could be determined with the R1 datasets, probably because of the very low number of genomic rNMPs in *H. volcanii* strains. On the other hand, no interception between red and blue curves for *T. barophilus* strains was found in the R1 and R2 datasets (Figure 3B), consistent with low origin utilization in actively proliferating *T. barophilus* cell lines (73).

**Figure 3.**
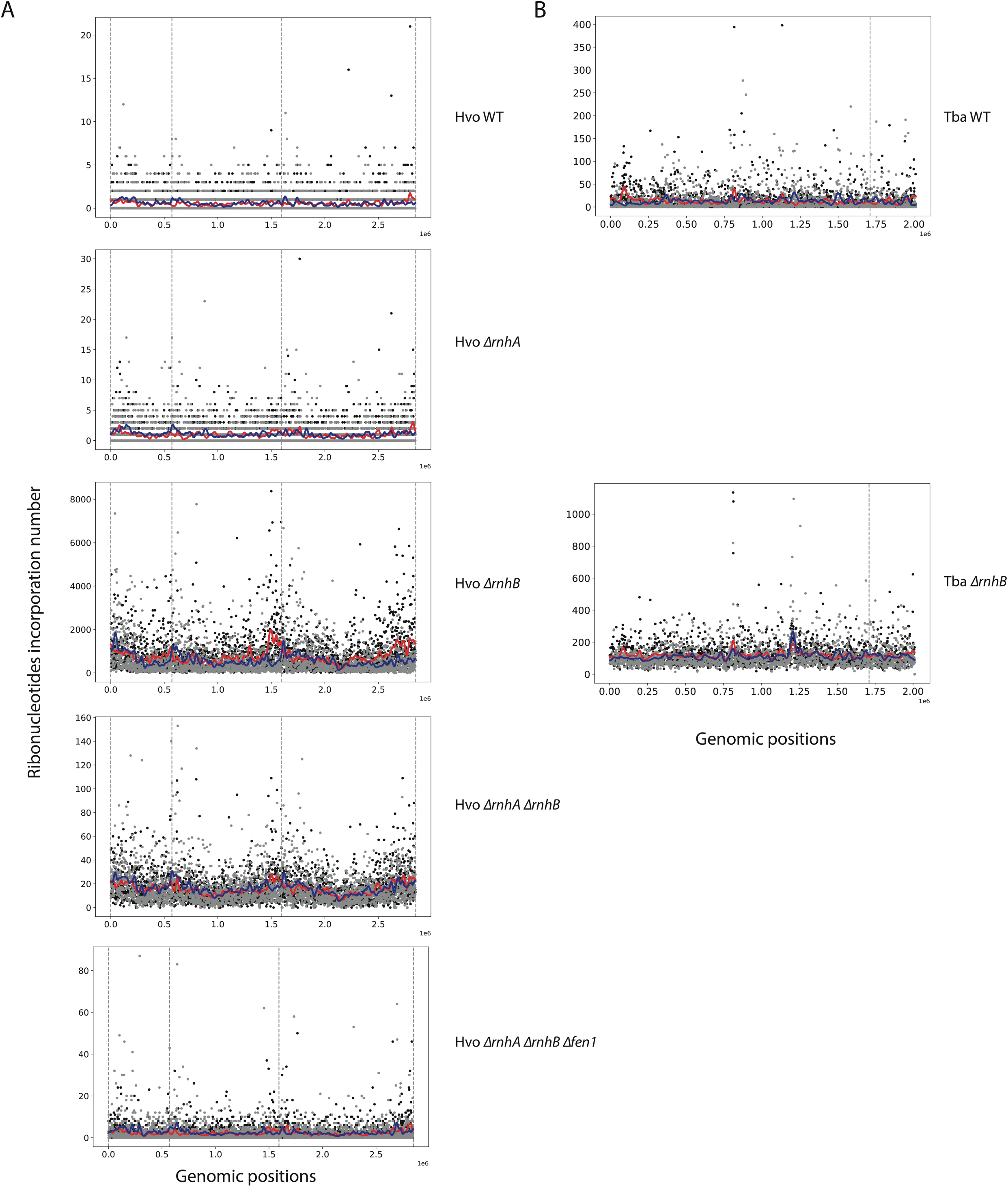
Chromosome-wide profiles of strand-specific ribonucleotide incorporation. **(A)** Chromosome of Hvo strains. **(B)** Chromosome of Tba strains. Locations and rNMP occurrences are plotted as dots on the graph. Black dots represent rNMP incorporation on the forward strand, while grey dots are rNMP incorporation on reverse strands. Red and Blue curves are obtained after application of one-dimensional Gaussian filter to the forward and reverse strand-containing embedded rNMPs. Vertical dashed-lines give the position of the *Ori* sites. R2 datasets are shown.

### Changing rNMP-base composition in the genome of *ΔrnhB* mutant cell lines

Considering the identity of the rNMPs incorporated in genomic DNA of wild-type and RNase H mutants for each library (Hvo and Tba), a shift in their distribution were clearly observed upon deletion of RNase HII (compare wild-type and Δ*rnhB* containing*-*mutant libraries in R1 and R2 datasets) (Figure 4, Table 2 and Table S5). For instance, rC and rU were preferred in the DNA of Hvo WT for R1 datasets (FS101 library) in comparison to rA and to a lesser extent rG in *ΔrnhB* mutant cell lines (FS100 and FS102 libraries) (Figure 4A and Table S5). However, in R2 datasets, a marked preference for rG in Hvo WT (FS220 library) was shifted toward rC and rA along with a strong reduction in rU level particularly in *ΔrnhB* mutant cell lines (FS218, FS219 and FS223 libraries) (Figure 4A and Table 2). No major change in the rNMP-base composition was observed in *ΔrnhA* mutant as compared to wild-type libraries (FS217 and FS220, R2 dataset, respectively) (Figure 4 and Table 2). Analysis of heatmap frequencies of each rNMP in the genome of Tba *ΔrnhB* shows an increased level of rC along with a clear reduction in rU in both R1 and R2 datasets (FS99 and FS221 libraries, respectively) (Figure 4B, Table 2 and Table S5) in comparison with rC and rG preference for their respective wild-type libraries in R1 and R2 datasets (FS98 and FS222)(Figure 4B, Table 2 and Table S5).

**Figure 4.**
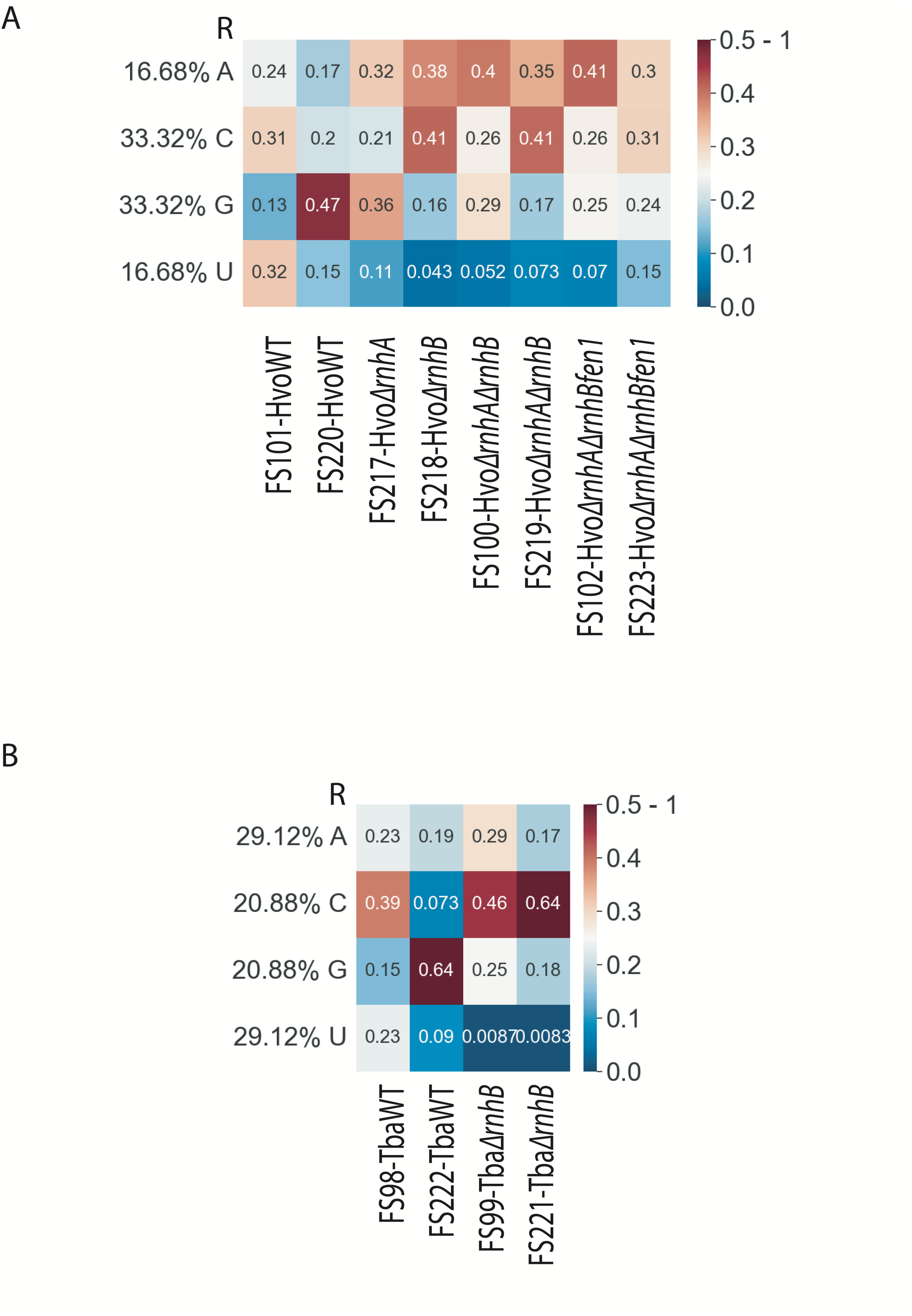
rNMP-base frequencies in the genome of wild-type and mutant *H. volcanii* and *T. barophilus* cells. The relative frequency of ribonucleotide incorporation normalized to the frequency of the corresponding base in the reference genome of all libraries is shown. **(A)** *H. volcanii* genotypes. **(B)** *T. barophilus* genotypes. Each row corresponds to one rNMP-base and each column represents one ribose-seq library. The heatmap colour scale of frequencies values is on the right with blue to white for 0 to 0.25 and white to brown for 0.25 to 0.5-1.

**Table 2.** Rate of rNMP incorporation (%) for the R2 datasets. (N/A stands for data missing after the ribose-map process).

| Species | Genotypes | Genetic element | rA | rC | rG | rU |
| --- | --- | --- | --- | --- | --- | --- |
| Hvo | WT | Chr | 16.9 | 20.5 | **47.7** | 14.9 |
|  |  | pHV1 | 12.2 | 14.2 | **61.4** | 12.2 |
|  |  | pHV3 | 18.1 | 20.6 | **43.9** | 17.3 |
|  |  | pHV4 | 16.7 | 22.7 | **46.3** | 14.4 |
| | $\Delta rnhA$ | Chr | 31.1 | 21.6 | **36.2** | 11.1 |
|  |  | pHV1 | 27.8 | 23.3 | **39.6** | 9.2 |
|  |  | pHV3 | 31.5 | 23.0 | **34.8** | 10.7 |
|  |  | pHV4 | 32 | 21.6 | **34.1** | 12.3 |
| | $\Delta rnhB$ | Chr | 37.4 | **41.9** | 16.5 | 4.3 |
|  |  | pHV1 | 37.1 | **40.5** | 18.4 | 4.0 |
|  |  | pHV3 | 38.5 | **41.6** | 16.0 | 3.9 |
|  |  | pHV4 | 37.0 | **42.1** | 17.2 | 3.8 |
| | $\Delta rnhA \Delta rnhB$ | Chr | 34.2 | **41.5** | 17.1 | 7.2 |
|  |  | pHV1 | 35.9 | **39.1** | 18.1 | 6.9 |
|  |  | pHV3 | 35.1 | **40.9** | 16.5 | 7.4 |
|  |  | pHV4 | 34.3 | **41.5** | 17.4 | 6.8 |
| | $\Delta rnhA \Delta rnhB \Delta fen1$ | Chr | 29.9 | **31.2** | 24.2 | 14.6 |
|  |  | pHV1 | N/A | N/A | N/A | N/A |
|  |  | pHV3 | 33.3 | **32.3** | 21.8 | 12.7 |
|  |  | pHV4 | 28.0 | **32.6** | 27.0 | 12.4 |
| Tba | WT | Chr | 19.4 | 7.3 | **64.2** | 9 |
| | $\Delta rnhB$ | Chr | 17.6 | **63.9** | 17.6 | 0.8 |

### Flanking base distribution at rNMP incorporation sites

We then analysed the sequence context and base composition surrounding the incorporation sites of rNMPs (Figure 5). The base distribution is depicted in two graphical representations, one that comprises all the genomic coordinates without distinction of the ribonucleotide bases (Figure 5, “Combined” plots) and the other considering the identity of each ribonucleotide base (Figure 5, rA, rC, rG and rU plots).

**Figure 5.**
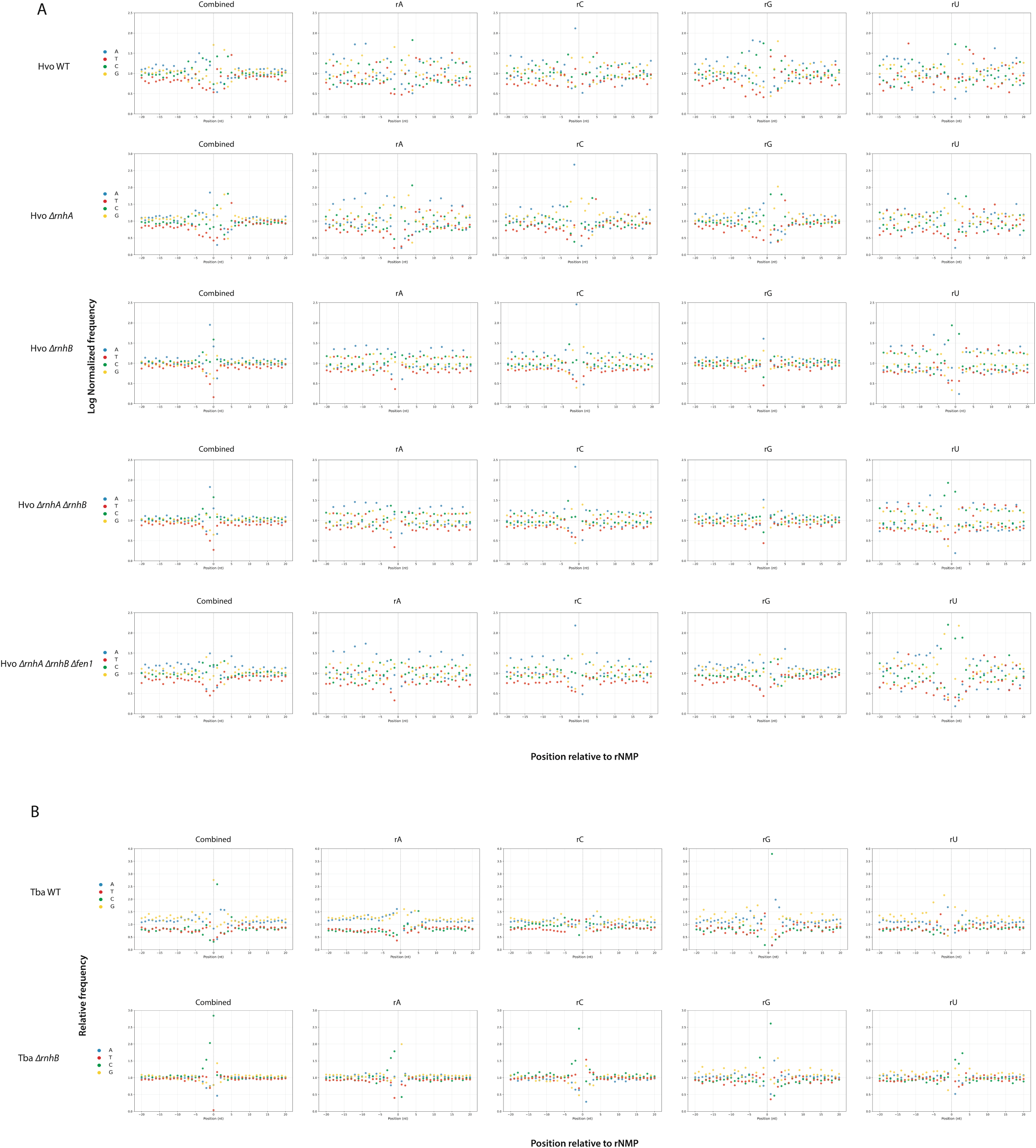
Distribution of DNA base sequences at rNMP incorporation sites. **(A)** Chromosome of Hvo strains. **(B)** Chromosome of Tba strains. For each strain, the base distribution encountered at the genomic coordinate of the rNMP incorporation site is computed for all four rNMP (“combined”) and separately for each rNMP (rA, rC, rG and rU). Position 0 on the x-axis represents the site of rNMP incorporation, − and + positions represent upstream and downstream dNMPs, respectively. The y-axis shows the frequency of each base found in the genomic context surrounding the rNMP incorporation sites normalized to the frequency of the corresponding base in the reference genome. R2 datasets are shown.

For *H. volcanii* strains, three patterns were noticeable in the chromosomes (Figure 5A, “Combined” plots, left part, R2 datasets). First, a 5-base sequence (5’-CGGCT-3)’ was found downstream the rNMP site in Hvo WT and *ΔrnhA* (Figure 5A, “Combined” plots, left part). Then, a single-base sequence involving base A was detected at the −1 position not only in the chromosome of *ΔrnhA*, but also in that of Hvo *ΔrnhB*, Hvo *ΔrnhAΔrnhB* and Hvo Δ*rnhA*Δ*rnhB*Δ*fen1* (blue dot in Figure 5A, “Combined” plots, left part). Finally, another single-base A was predominantly found at +5 position relative to the rNMP sites in Hvo Δ*rnhA*Δ*rnhB*Δ*fen1* (blue dot in Figure 5A, “Combined” plots, left part). When the base distribution was computed separately according to each nucleotide (Figure 5A, rA, rC, rG or rU plots), the 5-base sequence (5’-CGGCT-3’) slightly changed for rC in Hvo *ΔrnhA* and Hvo WT strains (Figure 5A, compare combined and rC plots), in which base G replaced base C at +1 position relative to the incorporation site. Moreover, a single base-sequence pattern involving base A at position −1 was strikingly conserved for rC compared to the three other ribonucleotides in all mutant Hvo strains, excepted for Hvo *ΔrnhB* (Figure 5A, compare rC plots with rA, rG or rU plots). Consistently, these patterns surrounding the rNMP sites in the chromosome in all Hvo strains were similarly conserved in the genetic elements of Hvo (Figures S3A, S4A and S5A), indicating that the slight difference in their GC content is not influencing (Table S1). Comparatively to R2 datasets, distinct patterns of flanking base distribution relative to the rNMP sites were found in the chromosome all Hvo strains in R1 datasets with a high degree of similarity between the mutated strains for all and each ribonucleotide base (“Combined” plots and rA, rC, rG and rU plots compared between Figure 5 and Figure S6A). It should be noted that fuzzy patterns observed in wild-type Hvo libraries for R1 datasets were likely caused by low number of rNMP occurrences (Figure S6A).

For *T. barophilus*, a clear difference of flanking base distribution between WT and *ΔrnhB* strains was observed (Figure 5B, “Combined” plots, left part, R2 datasets). While a 5-base sequence of 5’-CAAGC-3’ was predominant upstream of the rNMP sites in the chromosome of Tba WT, a 3-base pattern of 5’-CCC-3’ downstream of the incorporation sites and base G at +1 position were found in the chromosome Tba *ΔrnhB* (Figure 5B, “Combined” plots, left part). When base distribution was analysed separately for each ribonucleotide base, different patterns were observed (Figure 5B, rA, rC, rG and rU plots). In Tba WT, a 5-base sequence pattern of 5’-GGAGC-3’ was found for rA and rU (Figure 5B, rA and rU plots), whereas only bases T and C were detected at +1 position relative to rC and rG (Figure 5B, rC and rG plots, respectively). In Tba *ΔrnhB*, the 5’-CCC-3’ pattern occurred only for ribonucleotides rA and rC at the −1 position relative to the rNMP site (Figure 5B, compared rA, rC and rG, rU plots). Comparatively to R2 datasets, distinct base distributions were found in Tba WT R1 datasets (Figure S6B). For Tba WT when the identity of the nucleotide was ignored (Figure S6B, combined plot, left part), rNMP incorporation site was flanked by single base A at −1 position and single base T at +1 position. This pattern was also found for rA and rU (Figure S6B, rA and rU plots) whereas only base A at −1 position was present for rG (Figure S6B, rG plot) and only base T at +1 position for rC (Figure S6B, rC plot). For Tba *ΔrnhB*, rNMP incorporation site is flanked by base C at −1 position and base G at +1 position (Figure S6B, combined plot, left part). This pattern was also found for rA (Figure S6B, rA plot) whereas only base C at −1 position was strongly preferred for rC (Figure S6B, rC plot), base G surrounded rG (−1 and +1 positions)(Figure S6B, rG plot) and a pattern of four bases C was clearly observed downstream of rU (Figure S6B, rU plot). Overall, these results, demonstrated the influence of nucleotide environment on rNMP embedment regards to RNase H genotypes. They also showed that the different sets of restriction enzymes used during the library preparation protocols did account for rNMP sequence context variations.

### Frequencies of downstream and upstream deoxyribonucleotide base at embedded rNMPs

To get a narrow view of the surrounding dNMPs at the rNMP incorporation sites, the exact frequencies of dNMPs (N) immediately upstream (at −1 position) or downstream (at +1 position) of the embedded rNMPs (R) were calculated and plotted as dinucleotide-heatmaps (16 dinucleotide combinations). It is of note that the different restrictions enzyme sets used in the libraries did account for variations in the dinucleotide frequencies between R1 and R2 datatsets (Figure 6, left and right panels, respectively).

**Figure 6.**
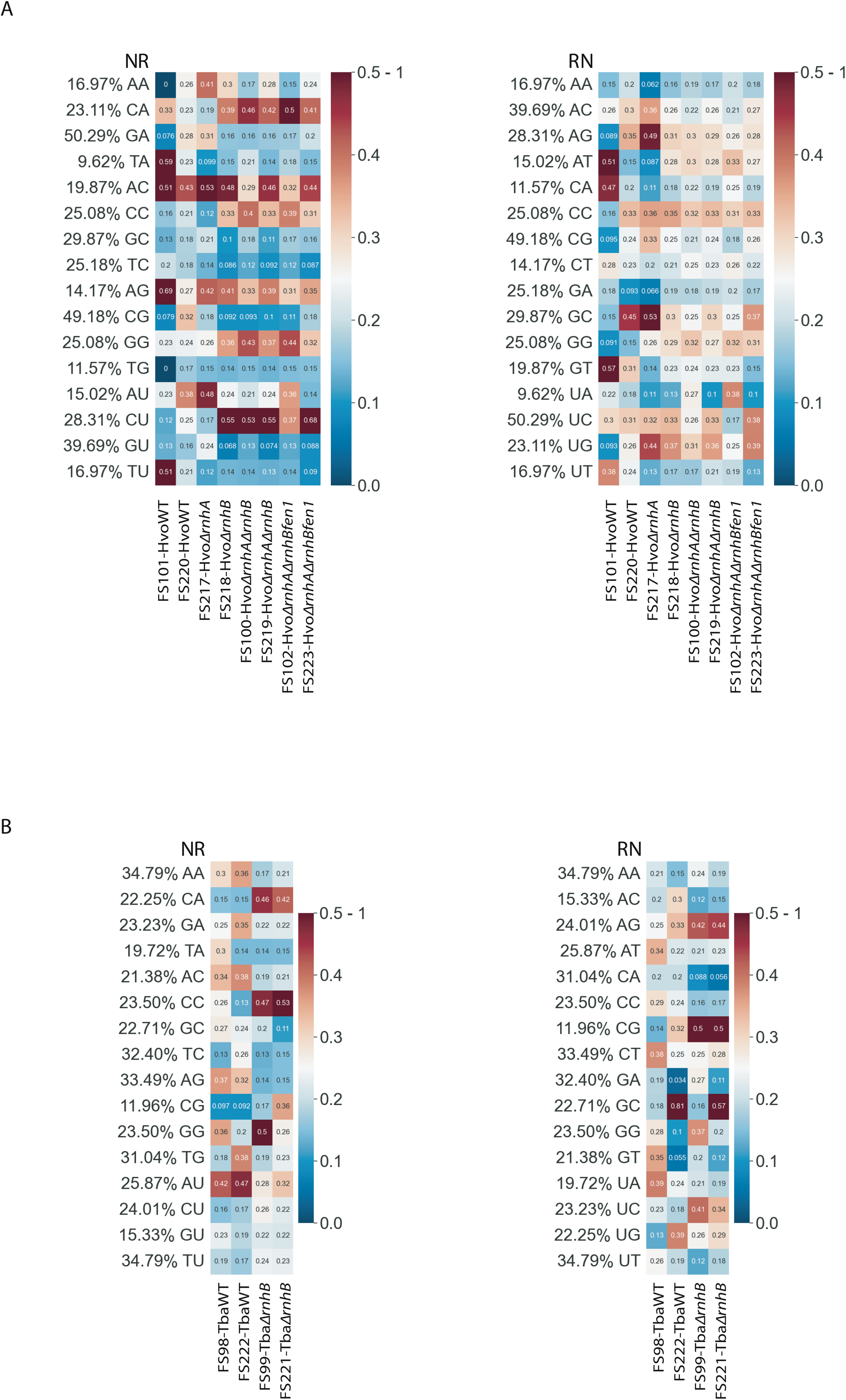
Heatmaps of dinucleotide normalized frequencies for *H. volcanii* (A) and *T. barophilus* (B). For each rNMP (R), the dNMP (N) directly upstream (left panels) and the dNMP directly downstream (right panel) were determined and the normalized frequencies were calculated (See Methods section for the formula). Each row corresponds to one combination of dinucleotides (NR or RN) and each column represents one ribose-seq library. The heatmap colour scale of frequencies values is on the right with blue to white for 0 to 0.25 and white to brown for 0.5-1.

Analysis of the NR dinucleotide-heatmap in *H. volcanii* strains demonstrated preferences for dCMP: CrA, CrC, and, particularly, CrU, and for dAMP: ArC and ArG, and for GrG in all Δ*rnhB* as compared to wild-type and Δ*rnhA* libraries, in which we observed a conserved high preference for ArC (Figure 6A, left, NR dinucleotide combinations). In comparison to FS101-HvoWT that showed marked NR-dinucleotide preferences for dTMP: TrA and TrU, and for dAMP: ArC and ArG, FS220-HvoWT did not have strong preferences for NR except ArC. For Hvo *ΔrnhA*, NR-dinucleotide preferences for dAMP: ArA, ArU, ArC and ArG were mainly observed, in which ArC and ArG were conserved in all Δ*rnhB* libraries (Figure 6A, left, NR dinucleotide combinations). Considering RN dinucleotide combinations with its downstream dNMP (+1 position regards to rNMP site)(Figure 6A, right panel, RN dinucleotide combinations), weak but conserved preferences for dGMP: rAG, rGG and rUG, and for dCMP: rGC, rCC, and for rAT were observed in all Δ*rnhB* as compared to wild-type and Δ*rnhA* libraries (Figure 6A, right, RN dinucleotide combinations). For Hvo *ΔrnhA*, stronger NR-dinucleotide preferences for rAG, rGC, and rUG were detected as compared to all Δ*rnhB* libraries. The dinucleotide rGC was mainly preferred in FS220-HvoWT, whereas FS101-HvoWT showed marked RN-dinucleotide preferences for rAT, rCA and rGT.

Analysis of the NR dinucleotide-heatmap in *T. barophilus* strains showed a strong preference for dCMP: CrA and CrC in Δ*rnhB*, as compared to wild-type libraries (Figure 6B, left, NR dinucleotide combinations). For the wild-type libraries (FS98-TbaWT and FS222-TbaWT), we did notice weak but conserved preferences for dAMP: ArA, ArC, ArG and ArU. In the heatmap of RN-dinucleotide combinations, there was the same and marked preference for dGMP: rAG and rCG in Δ*rnhB*, as compared to wild-type libraries (Figure 6B, right, RN dinucleotide combinations). Dissimilarities in the RN dinucleotide combinations were mainly seen in wild-type Tba libraries, in which the dinucleotide rGC had only the strongest frequencies in FS222-TbaWT.

### Nucleotide pool quantification regardless of RER dysfunction and ribonucleotide incorporation

Because our genome-wide analyses of rNMP incorporation highlighted specific features of incorporation in wild-type and RNase H-deficient cells, we thus questioned about possible influence of the level and balance of NTPs and dNTPs. This is of particular interest since recent studies have shown a direct link between deoxyribonucleoside triphosphate/ribonucleoside triphosphate (dNTP/rNTP) ratios and the frequency of rNMP incorporation in DNA (56–58). Here we conducted dNTP and rNTP measurements in wild-type and RNase H mutant cells of both species. The levels of dNTPs and rNTPs were evaluated from exponentially growing cells according to the recently optimized UV-based HPLC protocol which allows the simultaneous detection of dNTPs, rNTPs and ADP (68). The obtained ATP/ADP ratios were satisfying enough to validate our measurements (see Tables in Figure 7)(74).

**Figure 7.**
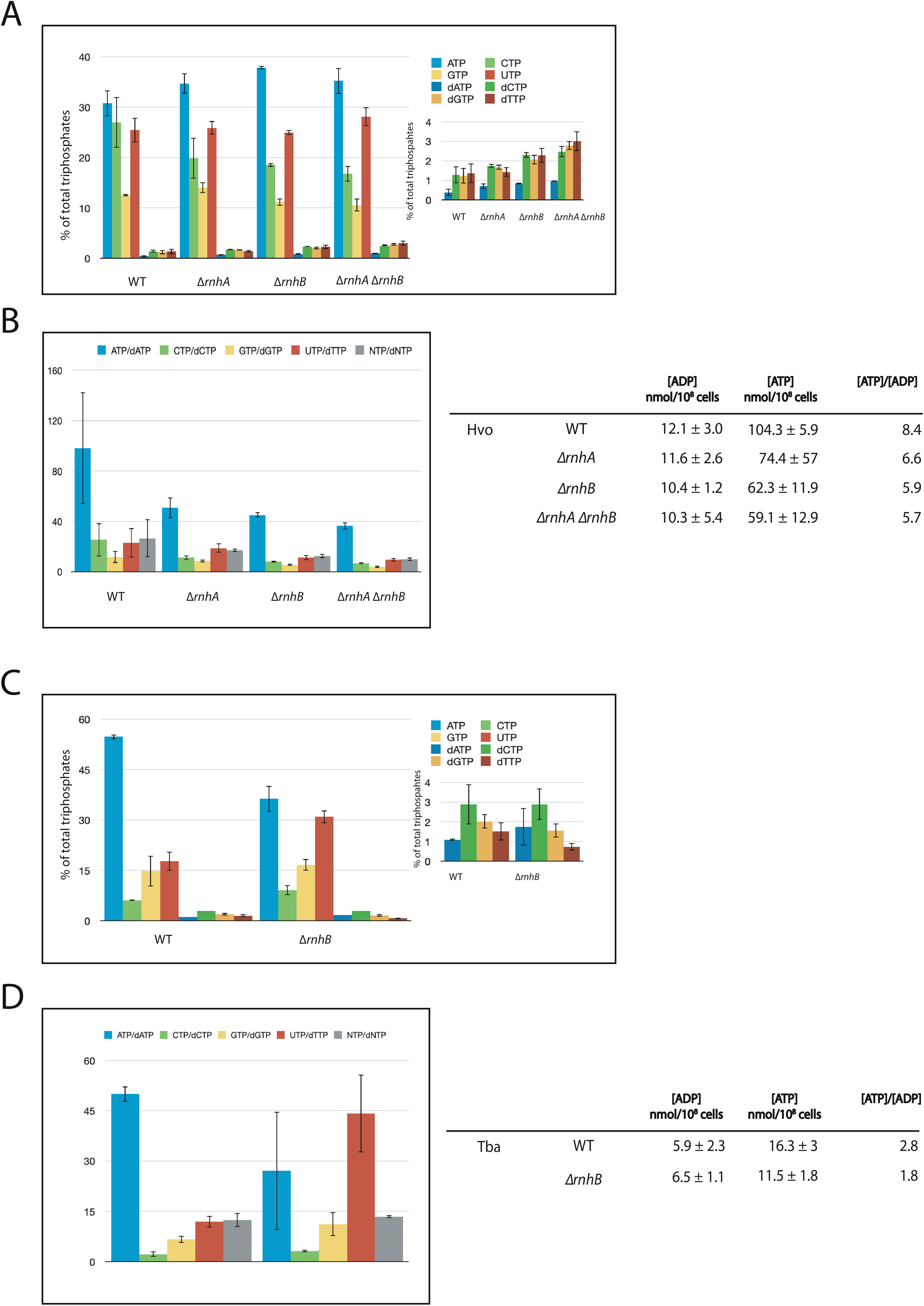
Intracellular levels of dNTPs and rNTPs in wild-type and mutant RNase H dividing cells. **(A)** NTPs and dNTPs in wild-type and RNase H null *H. volcanii* cells. The upper chart contains the percentage of the total triphosphate pool (*i.e.* ATP, CTP, GTP, UTP, dCTP, dGTP, dTTP and dATP). The inset shows a magnification of the dNTP pools. **(B)** The results in A are replotted as the ratio of each NTP to the corresponding dNTP in B, ATP/dATP, CTP/dCTP, GTP/dGTP, UTP/dTTP and total NTP/dNTP ratio values of the Hvo strains. The table on the right presents the ADP and ATP concentration values and their ratio. **(C)** NTPs and dNTPs in wild-type and RNase H null *T. barophilus* cells. The upper chart contains the percentage of the total triphosphate pool (*i.e.* ATP, CTP, GTP, UTP, dCTP, dGTP, dTTP and dATP). The inset shows a magnification of the dNTP pools. **(D)** The results in C are replotted as the ratio of each NTP to the corresponding dNTP in C, ATP/dATP, CTP/dCTP, GTP/dGTP, UTP/dTTP and total NTP/dNTP ratio values of the Tba strains. The table on the right presents the ADP and ATP concentration values and their ratio.

In *H. volcanii* cells, the results demonstrated that ATP was the most abundant ribonucleoside triphosphate for all while GTP was the lowest (Figure 7A, blue and yellow bars, respectively). The relative quantities of dNTPs remained unchanged (dCTP≃dGTP≃dTTP>dATP) (Figure 7A, right inset) while those of rNTPs differed between wild-type (ATP≃CTP≃UTP>GTP) and RNase H mutants (ATP>UTP>CTP>GTP) (Figure 7A). It is interesting to note that the proportion of dNTPs slightly increased in RNase H mutants compared to the wild-type *H. volcanii* strain, with Hvo *ΔrnhAΔrnhB* showing the greatest augmentation (Figure 7A, right inset). On the other hand, the total rNTP level was 23-fold, 16-fold, 12-fold or 10-fold higher than total dNTP content for Hvo WT, Hvo *ΔrnhA*, Hvo *ΔrnhB* and Hvo *ΔrnhAΔrnhB*, respectively, (Figure 7B, gray bars), corroborating previous data showing the excess of rNTPs over dNTPs in archaeal, bacterial and eukaryotic cells (4,5,58,75,76). According to the identity of the base, the ratio of ATP/dATP was higher than all other NTP/dNTP ratios (UTP/dTTP, CTP/dCTP, and GTP/dGTP) in all *H. volcanii* strains (Figure 7B), while being diminished in RNase H mutant compared to wild-type cells (Figure 7B, blue bars).

In *T. barophilus* strains, comparison of relative quantities of dNTPs showed a different balance between Tba WT (dCTP>dGTP>dTTP>dATP) and Tba *ΔrnhB* (dCTP>dATP≃dGTP>dTTP) (Figure 7C, right inset), whereas those of rNTPs were of the same order (ATP>UTP>GTP>CTP) (Figure 7C). Nonetheless, a decrease of ATP proportion (∼1.6-fold) and an increase of UTP level (∼2-fold) were clearly noticeable in Tba *ΔrnhB* compared to Tba WT (Figure 7C, blue and red bars, respectively). Moreover, an increase of dATP level and a decrease of dTTP level were observable in Tba *ΔrnhB* compared to Tba WT (Figure 7C, blue and red bars in right inset, respectively). The total rNTP content was 13-fold higher than the total dNTP content for Tba WT and Tba *ΔrnhB* (Figure 7D, grey bars). According to the identity of the base, the ATP/dATP was higher than all other NTP/dNTP (UTP/dTTP, CTP/dCTP, and GTP/dGTP) in all *T. barophilus* strains (Figure 7D), while being reduced in RNase HII mutant cells (Figure 7D, blue bars).

## DISCUSSION

The recent availability of methods allowing the genome-wide mapping of the embedded rNMP in DNA has given a new impetus to the characterization of this process (49,50,52–55). It gives the possibility to confirm biochemical *in vitro* results of rNMP misincorporation and contributes to the comprehensive view of *in vivo* cellular mechanisms including the associated physiological consequences. In this study, we evaluated the effect of several genotypes from two archaeal species on rNMP genomic embedding. The results confirm the non-random character of rNMP incorporation and the prominent role of type 2 ribonuclease H in genomic homeostasis as previously revealed in other domains of life. In *H. volcanii*, persistent rNMPs in the genome of WT and *ΔrnhA* cells display comparable patterns of incorporation, which differs to that of *ΔrnhB* cells. This result is similar to the ones obtained in the nuclear DNA of *Saccharomyces cerevisiae*, where identical patterns were observed between WT and *rhn1-null* cells but not with *rhn201-null* cells (57). Based on *in vitro* evidences, different RNase H processing of RNA/DNA heteroduplexes have been described, in which type 2 RNase H cuts all types of RNA/DNA hybrids including single rNMPs while type 1 RNase H cleaves DNA heteroduplexes with at least four consecutive embedded rNMPs (77). Moreover, type 1 RNase H can hydrolyse duplexes with three internal rNMPs, but at a slower catalysis (40). Based on our differential results, we propose that the two types of RNase H in *H. volcanii* may operate on different substrates *in vivo*, with the assumption that stretches of rNMPs (≥4 rNMPs) might be less prominent than single rNMPs in dsDNA. It is of note that the propensity of single *versus* covalently linked ribonucleotides in genomic DNA has not been methodologically resolved. To date, *in vitro* RNase H fragmentation of plasmids from *E. coli* and genomic DNA from mouse cells are the main studies accounting for a lower density of rNMP stretches than single or, at most, two internal rNMPs embedded in genomic DNA (29,37). On the other hand, *T. barophilus* evolved a single type 2 RNase H that clearly alleviates embedded rNMPs in the chromosome. However, the balance between single or longer stretches of rNMPs still remains unclear. Examination of the global distribution of rNMP sites across the genomes did not reveal particular hotspots in the genome of the wild-type and mutant *H. volcanii* strains but rather the presence of coldspots which apparently locate into the 16S and 23S ribosomal RNA genes. In contrast, hotspots were mainly detectable in the chromosome of *T. barophilus* and clustered in 16S ribosomal RNA gene. It is possible that actively transcribed ribosomal RNA genes in these two *Archaea* are organized in different chromatin landscapes, which facilitates the nascent RNA to hybridise (or not) back to the DNA template. As such, the presence of rNMPs may reflect imperfect removal of DNA-RNA hybrids in the genomes, which may occur in the genome of exponentially growing *T. barophilus* cells, but not in *H. volcanii*. A recent genomic study demonstrated the absence of correlation between chromatin accessibility and transcriptional activity in *H. volcanii*, likely being refractory to hybridization of nascent RNA and precluding any accumulation of persistent rNMPs (78). Although not yet documented in *T. barophilus*, it would be thus interesting to analyse chromatin landscape in order to correlate or not with the formation of transient RNA-DNA hybrids.

Ribose-seq strand-specificity analysis shows a slight preferential rNMP segregation on the reverse strand in the chromosome, but mainly on the forward strand of most plasmids of all *H. volcanii* strains (Table S3). These chromosome-based features were not influenced by the absence of RNases H and neither correlated with GC content or GC skew profile. Nearly equal on both strands of wild-type *T. barophilus*, rNMP enrichment was rather detectable on the forward strand of RNase HII mutant. Nevertheless, it remains unclear whether this genotype-correlation arises because of genetic variation or/and is related to the archaeal cell lineage. *H. volcanii* and *T. barophilus* cells are two archaeal species with genomic divergences and particular lifestyles, which may account for the discrepancy. Examining the frequency of strand-specific rNMPs across the chromosomes reveals a strong asymmetry of rNMP segregation in *H. volcanii* cells (Figure 3), whose curves intercept at chromosomal replication origins *oriC1*, *oriC2* and *oriC3* (0 kb, 571 kb and 1593 kb, respectively). Only detectable in RNase HII mutants, this oriC-intersecting profile is coincident with origin firing (72). Conversely, no asymmetric pattern is apparent in exponentially growing *T. barophilus* cells, consistent with low origin utilization in actively proliferating *T. barophilus* cell lines (73). This strand-specific transition at origins was previously described in *E. coli* and yeast by using three different methods for mapping genomic ribonucleotides (49,53,54), and particularly with replicase variants promiscuous for ribonucleotide incorporation in RER-deficient strains (*ΔrnhB)*. Here, strand switches at origins were detectable in *H. volcanii* cells lacking only the RNase HII enzyme, suggesting the natural propensity of replicases to incorporate rNMPs during the log phase of growth. Consistently, family D DNA polymerase in Euryarchaeota has been shown to embed rNMPs in DNA (4), likely due to its ancestral RNA polymerase catalytic core (79). Nevertheless, strand-specificity transition at *oriC* in replicating RER-deficient *T. barophilus* cells (*ΔrnhB)* was not visible, possibly because of low level of origin firing (73). Like for *E. coli* and yeast, it would be thus interesting to map genomic ribonucleotides in RER-deficient cells (*ΔrnhB) with* DNA polymerase variants (Family B, Family D or PriSL) promiscuous for rNMPs insertion. This would help to elucidate their *in vivo* role during DNA replication.

In this study, we also attempted to correlate the identity of rNMP incorporation with the varying pool of nucleotides in RER-proficient and RER-deficient archaeal cell lines. In general, rCMP is predominantly incorporated in the genome of archaeal cells lacking RNase HII, and its incorporation frequency coincides with a slight decrease of cellular CTP compared to wild-type cells in *H. volcanii*. However, no relationship is observed between the major genomic rGMP incorporation and the level of CTP in wild-type and RNase HI-null *H. volcanii* cells. Intriguingly, elevated level of UTP correlates with lower genomic rUMP in RNase HII-null *T. barophilus* cells. Nevertheless, our datasets do not allow assigning or/and predicting any genotype-correlated rNMPs regards to balancing nucleotide ratios. Further genetic constructions like crossing RNR (ribonucleotide reductase) and RNase H mutant strains are awaiting in order to evaluate the influence of varying dNTP/NTP ratios on genomic rNMP incorporation profiles of RER-deficient strains, as previously reported (56).

To go further in the analysis of rNMP retention in dsDNA, we examined the frequencies of downstream and upstream deoxyribonucleotide base at rNMP-embedment. The results demonstrate that specific preferences for the downstream and upstream deoxyribonucleotide bases surrounding rNMP sites are found in RNase HII-null cells, which might (or not) be shared with RNase HI-null cells, but not with wild-type *H. volcanii* cells. As highlighted, preference specificities also occur in the absence of RNase HII in *T. barophilus* cells. Different lines of evidences might explain these particular over-represented RN or NR dinucleotide combinations. First, there is the rNMP incorporation capacity of family D DNA polymerase (PolD) (4) which might be affected by the nature of the downstream and upstream template base, local sequence context as well as by primer length, rendering these particular patterns. In addition, RNA priming activity by primase/polymerase complex (PriSL) (80,81) might be influence by local sequence context such as specific recognition motif in the template, reflecting these favoured dinucleotide-rNMP motifs. Combined to specific and stable rNMP incorporations in dsDNA, the results also account for cross talk activities by type 1 and type 2 RNases H in wild-type *H. volcani* cells, and point out varying rNMP sites resistant to RNase H cleavage. Thus, the lack of either type 1 or type 2 RNases H allows to highlight the influence of surrounding base at the sites of rNMP incorporation reciprocally removed, in which the rate of RNase HII cleavage activity is the most predominant, while showing altered capacities in the presence of dCMP and dAMP upstream bases only. Downstream bases at rNMP sites do not seem to exert a strong effect on rNMP processing. In wild-type *T. barophilus* cells, variations of deoxynucleotide bases at rNMP sites are more frequent than for *H. volcanii*, but the lack of the single-RNase HII enzyme clearly reflects that both the upstream and downstream bases affect rNMP removal efficiency.

Being equipped like their bacterial and eukaryotic RNase HII/2 counterparts such as the four conserved carboxylates (DEDD motif) in the active site (Figure S7A-B and Figure S9B), the GRG 2’-OH sensing motif, the conserved tyrosine and residues involved in substrate binding (Figure S9B), it remains difficult to point out the molecular determinants involved in differential activity. Structural superimposition of archaeal RNase HII model predictions with the bacterial *T. maritima* RNase HII or the human RNase H2A catalytic subunit also demonstrates the high conservation of RNase HII/2 structure like the active site architecture and the so-called RNase H-fold (Figure S7B-C). Nevertheless, the overall structural similarity expressed as the RMSD over the Cα atoms of the whole protein assigns HvoRNase HII and TbaRNase HII rather to a functional of human RNase H2A than to *E. coli* RNase HII enzyme (Figure S7C), which is consistent with the higher sequence identities shared with the eukaryotic enzymes (Figure S9A), as previously noticed (82). Thus, we propose that the subtle structural variation in RNase HII archaeal enzymes may be involved in the variations of the identity of rNMP-bases and the surrounding deoxyribonucleotide bases in the archaeal genome.

It is also not excluded that other RNases H may operate in ribonucleotide repair, particularly, in *H. volcanii*. Comparatively to wild-type strains, single deletion *rnhA* mutant displays similar rNMP incorporation with rGMP the most representative. Although the *in vivo* role of the RNase HI under this study (Hvo_0732) is not yet fully understood, it is a good candidate for removing ribonucleotides in dsDNA. Harbouring the highest sequence identities (Figure S10A) and conserved catalytic amino acids (Figure S10B and Figure S8A-B) compared to the three others (Hvo_A0463, Hvo_A0277, and Hvo_2438), Hvo_0732 may act on rNMPs embedded in dsDNA. The conserved active site geometry along with the lowest RMSD value obtained with the bacterial counterpart (Figure S8B and Figure S8C) suggests plausible roles in eliminating single mismatched ribonucleotides in RNA stretches (25), in addition to RNA:DNA hybrid removal (RNA stretches, Okazaki fragments or R-loops) and strand-specific RER during replication(43). However, ribose-seq did not allow capturing these genomic RNA intermediates (52). Thus, in combination to RNase HII, Hvo_0732 RNase HI may also incise 3’ to single rNMP in dsDNA like the *E. coli* counterpart (42), which may explain the slight increased rNMP loads in ribose-seq and alkaline gel electrophoresis. Unexpectedly, the triple *rnhArnhBfen1* mutant displays lower rNMP incorporation frequency than the double *rnhArnhB* or the single *rnhB* mutant strains. Based on our structural superimposition with either bacterial or human RNase HI/1, Hvo_A0463 represents the most valuable substitute for RNase HI. Conservation of all four catalytic residues and active site architecture, with an additional N-terminal domain like that observed in Hvo_0732 (Figure S8A, B and C), suggest a possible back-up role of Hvo_A0463. For instance, ribonucleotide removal defective background would activate Hvo_A0463 *rnhA* gene expression which in turn takes over the function of missing RNases H. It will be of interest in future work to examine the level of gene expression at both protein and mRNA level in different RNase H genetic backgrounds, and whether these proteins functionally interact within a RER-complex or sequentially. *In vitro* specificity of RNases H1-like protein of *H. volcanii* also awaits further elucidation.

## ACKNOWLEDGEMENTS

The authors acknowledge the Pôle de Calcul et de Données Marines (PCDM; https://wwz.ifremer.fr/en/Research-Technology/Research-Infrastructures/Digital-infrastructures/Computation-Centre) for providing DATARMOR computing and storage resources. This work has received funding from the French Institute of Marine Research and Exploitation (IFREMER). The technical assistance of Josiane Lebars for preparation of *H. volcanii* cell culture medium throughout this project is greatly appreciated. The PhD student, M. Reveil, thanks ISblue, Interdisciplinary graduate school for the blue planet, for supporting her international mobility to Hofer’s laboratory.

## AUTHOR CONTRIBUTIONS

Yann Moalic: Conceptualization, methodology, investigation, validation, data curation related to high-throughput DNA sequencing work. Writing - review & editing the manuscript. Maurane Reveil: related to nucleotide pool quantification, conceptualization, methodology, investigation, validation, data curation, writing. Deepali L. Kundnani: related to all high-throughput DNA sequencing work, writing-review and editing ribose-seq data. Sathya Balachander: related to all high-throughput DNA sequencing work, ribose-seq methodology. Taehwan Yang: related to all high-throughput DNA sequencing work, ribose-seq methodology. Alli Gombolay: related to all high-throughput DNA sequencing work, ribose-seq methodology. Farahnaz Ranjbarian: related to nucleotide pool quantification, methodology. Raphael Brizard: related to *T. barophilus* cell culture, methodology. Patrick Durand: related to the establishment of nf-colabfold pipeline for protein structure predictions, conceptualization, methodology, data curation, writing. Hannu Myllykallio: related to *H. volcanii* cell culture, methodology, validation. Mohamed Jebbar: related to *T. barophilus* cell culture, methodology, validation. Anders Hofer: related to nucleotide pool quantification, conceptualization, methodology, investigation, validation, supervision, writing. Francesca Storici: related to all high-throughput DNA sequencing work, conceptualization, methodology, investigation, validation, supervision, writing. Ghislaine Henneke: Conceptualization, methodology, investigation, validation, data curation, supervision, funding acquisition, project administration, Writing - review & editing the manuscript.

## SUPPLEMENTARY DATA

Supplementary Data are available at NAR online.

## CONFLICT OF INTEREST

The authors declare no conflict of interest.

## FUNDING

M. R. was financially supported by Ifremer and the Brittany Regional Council. This work has received funding from the French Institute of Marine Research and Exploitation (Ifremer). Funding for open access charge : IFREMER.

## DATA AVAILABILITY

Data were submitted to NCBI under BioProject Accession Number : PRJNA1231464.

## Notes

### Competing Interest Statement

The authors have declared no competing interest.

## REFERENCES

1. McCulloch, S.D. and Kunkel, T.A. (2008) The fidelity of DNA synthesis by eukaryotic replicative and translesion synthesis polymerases. Cell Res, 18, 148–161.

2. Wang, W., Wu, E.Y., Hellinga, H.W. and Beese, L.S. (2012) Structural factors that determine selectivity of a high fidelity DNA polymerase for deoxy-, dideoxy-, and ribonucleotides. J Biol Chem, 287, 28215–28226.

3. Heider, M.R., Burkhart, B.W., Santangelo, T.J. and Gardner, A.F. (2017) Defining the RNaseH2 enzyme-initiated ribonucleotide excision repair pathway in *Archaea*. J Biol Chem, 292, 8835–8845.

4. Lemor, M., Kong, Z., Henry, E., Brizard, R., Laurent, S., Bosse, A. and Henneke, G. (2018) Differential Activities of DNA Polymerases in Processing Ribonucleotides during DNA Synthesis in *Archaea*. J Mol Biol, 430, 4908–4924.

5. Nick McElhinny, S.A., Watts, B.E., Kumar, D., Watt, D.L., Lundstrom, E.B., Burgers, P.M., Johansson, E., Chabes, A. and Kunkel, T.A. (2010) Abundant ribonucleotide incorporation into DNA by yeast replicative polymerases. Proc Natl Acad Sci U S A, 107, 4949–4954.

6. Yao, N.Y., Schroeder, J.W., Yurieva, O., Simmons, L.A. and O’Donnell, M.E. (2013) Cost of rNTP/dNTP pool imbalance at the replication fork. Proc Natl Acad Sci U S A, 110, 12942–12947.

7. Brown, J.A. and Suo, Z. (2011) Unlocking the sugar “steric gate” of DNA polymerases. Biochemistry, 50, 1135–1142.

8. Zatopek, K.M., Alpaslan, E., Evans, T.C., Sauguet, L. and Gardner, A.F. (2020) Novel ribonucleotide discrimination in the RNA polymerase-like two-barrel catalytic core of Family D DNA polymerases. Nucleic Acids Res, 48, 12204–12218.

9. Cilli, P., Minoprio, A., Bossa, C., Bignami, M. and Mazzei, F. (2015) Formation and Repair of Mismatches Containing Ribonucleotides and Oxidized Bases at Repeated DNA Sequences. J Biol Chem, 290, 26259–26269.

10. Jamsen, J.A., Sassa, A., Perera, L., Shock, D.D., Beard, W.A. and Wilson, S.H. (2021) Structural basis for proficient oxidized ribonucleotide insertion in double strand break repair. Nat Commun, 12, 5055.

11. Ordonez, H., Uson, M.L. and Shuman, S. (2014) Characterization of three mycobacterial DinB (DNA polymerase IV) paralogs highlights DinB2 as naturally adept at ribonucleotide incorporation. Nucleic Acids Res, 42, 11056–11070.

12. Nava, G.M., Grasso, L., Sertic, S., Pellicioli, A., Muzi Falconi, M. and Lazzaro, F. (2020) One, No One, and One Hundred Thousand: The Many Forms of Ribonucleotides in DNA. International Journal of Molecular Sciences, 21, 1706.

13. Ghodgaonkar, M.M., Lazzaro, F., Olivera-Pimentel, M., Artola-Boran, M., Cejka, P., Reijns, M.A., Jackson, A.P., Plevani, P., Muzi-Falconi, M. and Jiricny, J. (2013) Ribonucleotides misincorporated into DNA act as strand-discrimination signals in eukaryotic mismatch repair. Mol Cell, 50, 323–332.

14. Lujan, S.A., Williams, J.S., Clausen, A.R., Clark, A.B. and Kunkel, T.A. (2013) Ribonucleotides are signals for mismatch repair of leading-strand replication errors. Mol Cell, 50, 437–443.

15. Vengrova, S. and Dalgaard, J.Z. (2006) The wild-type *Schizosaccharomyces pombe* mat1 imprint consists of two ribonucleotides. EMBO Rep, 7, 59–65.

16. Sparks, J.L., Chon, H., Cerritelli, S.M., Kunkel, T.A., Johansson, E., Crouch, R.J. and Burgers, P.M. (2012) RNase H2-initiated ribonucleotide excision repair. Mol Cell, 47, 980–986.

17. Vaisman, A., McDonald, J.P., Noll, S., Huston, D., Loeb, G., Goodman, M.F. and Woodgate, R. (2014) Investigating the mechanisms of ribonucleotide excision repair in *Escherichia coli*. Mutat Res, 761, 21–33.

18. Reveil, M., Chapel, L., Vourc’h, B., Bosse, A., Vialle, L., Brizard, R., Moalic, Y., Jebbar, M. and Henneke, G. (2023) Processing of matched and mismatched rNMPs in DNA by archaeal ribonucleotide excision repair. iScience, 26, 108479.

19. Cerritelli, S.M., Iranzo, J., Sharma, S., Chabes, A., Crouch, R.J., Tollervey, D. and El Hage, A. (2020) High density of unrepaired genomic ribonucleotides leads to Topoisomerase 1-mediated severe growth defects in absence of ribonucleotide reductase. Nucleic Acids Res, 48, 4274–4297.

20. Sparks, J.L. and Burgers, P.M. (2015) Error-free and mutagenic processing of topoisomerase 1-provoked damage at genomic ribonucleotides. EMBO J, 34, 1259–1269.

21. Huang, S.N., Williams, J.S., Arana, M.E., Kunkel, T.A. and Pommier, Y. (2017) Topoisomerase I-mediated cleavage at unrepaired ribonucleotides generates DNA double-strand breaks. EMBO J, 36, 361–373.

22. Pommier, Y., Huang, S.-Y., Williams, J. and Kunkel, T. (2015) Topoisomerase-Induced DNA Cleavage at Ribonucleotide Misincorporation Sites. The FASEB Journal, 29, 371.373.

23. Williams, J.S., Smith, D.J., Marjavaara, L., Lujan, S.A., Chabes, A. and Kunkel, T.A. (2013) Topoisomerase 1-mediated removal of ribonucleotides from nascent leading-strand DNA. Mol Cell, 49, 1010–1015.

24. Malfatti, M.C., Balachander, S., Antoniali, G., Koh, K.D., Saint-Pierre, C., Gasparutto, D., Chon, H., Crouch, R.J., Storici, F. and Tell, G. (2017) Abasic and oxidized ribonucleotides embedded in DNA are processed by human APE1 and not by RNase H2. Nucleic Acids Res, 45, 11193–11212.

25. Shen, Y., Koh, K.D., Weiss, B. and Storici, F. (2012) Mispaired rNMPs in DNA are mutagenic and are targets of mismatch repair and RNases H. Nat Struct Mol Biol, 19, 98–104.

26. Vaisman, A., McDonald, J.P., Huston, D., Kuban, W., Liu, L., Van Houten, B. and Woodgate, R. (2013) Removal of misincorporated ribonucleotides from prokaryotic genomes: an unexpected role for nucleotide excision repair. PLoS Genet, 9, e1003878.

27. Nick McElhinny, S.A., Kumar, D., Clark, A.B., Watt, D.L., Watts, B.E., Lundstrom, E.B., Johansson, E., Chabes, A. and Kunkel, T.A. (2010) Genome instability due to ribonucleotide incorporation into DNA. Nat Chem Biol, 6, 774–781.

28. Lazzaro, F., Novarina, D., Amara, F., Watt, D.L., Stone, J.E., Costanzo, V., Burgers, P.M., Kunkel, T.A., Plevani, P. and Muzi-Falconi, M. (2012) RNase H and postreplication repair protect cells from ribonucleotides incorporated in DNA. Mol Cell, 45, 99–110.

29. Reijns, M.A., Rabe, B., Rigby, R.E., Mill, P., Astell, K.R., Lettice, L.A., Boyle, S., Leitch, A., Keighren, M., Kilanowski, F. et al. (2012) Enzymatic removal of ribonucleotides from DNA is essential for mammalian genome integrity and development. Cell, 149, 1008–1022.

30. Crow, Y.J., Leitch, A., Hayward, B.E., Garner, A., Parmar, R., Griffith, E., Ali, M., Semple, C., Aicardi, J., Babul-Hirji, R. et al. (2006) Mutations in genes encoding ribonuclease H2 subunits cause Aicardi-Goutieres syndrome and mimic congenital viral brain infection. Nat Genet, 38, 910–916.

31. Pizzi, S., Sertic, S., Orcesi, S., Cereda, C., Bianchi, M., Jackson, A.P., Lazzaro, F., Plevani, P. and Muzi-Falconi, M. (2015) Reduction of hRNase H2 activity in Aicardi-Goutieres syndrome cells leads to replication stress and genome instability. Hum Mol Genet, 24, 649–658.

32. Gunther, C., Kind, B., Reijns, M.A., Berndt, N., Martinez-Bueno, M., Wolf, C., Tungler, V., Chara, O., Lee, Y.A., Hubner, N. et al. (2015) Defective removal of ribonucleotides from DNA promotes systemic autoimmunity. J Clin Invest, 125, 413–424.

33. Aden, K., Bartsch, K., Dahl, J., Reijns, M.A.M., Esser, D., Sheibani-Tezerji, R., Sinha, A., Wottawa, F., Ito, G., Mishra, N. et al. (2019) Epithelial RNase H2 Maintains Genome Integrity and Prevents Intestinal Tumorigenesis in Mice. Gastroenterology, 156, 145–159 e119.

34. Hiller, B., Achleitner, M., Glage, S., Naumann, R., Behrendt, R. and Roers, A. (2012) Mammalian RNase H2 removes ribonucleotides from DNA to maintain genome integrity. J Exp Med, 209, 1419–1426.

35. Uehara, R., Cerritelli, S.M., Hasin, N., Sakhuja, K., London, M., Iranzo, J., Chon, H., Grinberg, A. and Crouch, R.J. (2018) Two RNase H2 Mutants with Differential rNMP Processing Activity Reveal a Threshold of Ribonucleotide Tolerance for Embryonic Development. Cell Rep, 25, 1135–1145 e1135.

36. Schroeder, J.W., Randall, J.R., Hirst, W.G., O’Donnell, M.E. and Simmons, L.A. (2017) Mutagenic cost of ribonucleotides in bacterial DNA. Proc Natl Acad Sci U S A, 114, 11733–11738.

37. Kouzminova, E.A., Kadyrov, F.F. and Kuzminov, A. (2017) RNase HII Saves rnhA Mutant *Escherichia coli* from R-Loop-Associated Chromosomal Fragmentation. J Mol Biol, 429, 2873–2894.

38. Lockhart, A., Pires, V.B., Bento, F., Kellner, V., Luke-Glaser, S., Yakoub, G., Ulrich, H.D. and Luke, B. (2019) RNase H1 and H2 Are Differentially Regulated to Process RNA-DNA Hybrids. Cell Rep, 29, 2890–2900 e2895.

39. Cerritelli, S.M. and Crouch, R.J. (2009) Ribonuclease H: the enzymes in eukaryotes. FEBS J, 276, 1494–1505.

40. Hogrefe, H.H., Hogrefe, R.I., Walder, R.Y. and Walder, J.A. (1990) Kinetic analysis of *Escherichia coli* RNase H using DNA-RNA-DNA/DNA substrates. J Biol Chem, 265, 5561–5566.

41. Ohtani, N., Tomita, M. and Itaya, M. (2008) Junction ribonuclease activity specified in RNases HII/2. FEBS J, 275, 5444–5455.

42. Tannous, E., Kanaya, E. and Kanaya, S. (2015) Role of RNase H1 in DNA repair: removal of single ribonucleotide misincorporated into DNA in collaboration with RNase H2. Sci Rep, 5, 9969.

43. Lazowski, K., Faraz, M., Vaisman, A., Ashton, N.W., Jonczyk, P., Fijalkowska, I.J., Clausen, A.R., Woodgate, R. and Makiela-Dzbenska, K. (2023) Strand specificity of ribonucleotide excision repair in *Escherichia coli*. Nucleic Acids Res, 51, 1766–1782.

44. Maduike, N.Z., Tehranchi, A.K., Wang, J.D. and Kreuzer, K.N. (2014) Replication of the *Escherichia coli* chromosome in RNase HI-deficient cells: multiple initiation regions and fork dynamics. Mol Microbiol, 91, 39–56.

45. Kouzminova, E.A. and Kuzminov, A. (2021) Ultraviolet-induced RNA:DNA hybrids interfere with chromosomal DNA synthesis. Nucleic Acids Res, 49, 3888–3906.

46. Cerritelli, S.M., Frolova, E.G., Feng, C., Grinberg, A., Love, P.E. and Crouch, R.J. (2003) Failure to produce mitochondrial DNA results in embryonic lethality in Rnaseh1 null mice. Mol Cell, 11, 807–815.

47. Manini, A., Caporali, L., Meneri, M., Zanotti, S., Piga, D., Arena, I.G., Corti, S., Toscano, A., Comi, G.P., Musumeci, O. et al. (2022) Case Report: Rare Homozygous RNASEH1 Mutations Associated With Adult-Onset Mitochondrial Encephalomyopathy and Multiple Mitochondrial DNA Deletions. Front Genet, 13, 906667.

48. Reyes, A., Melchionda, L., Nasca, A., Carrara, F., Lamantea, E., Zanolini, A., Lamperti, C., Fang, M., Zhang, J., Ronchi, D. et al. (2015) RNASEH1 Mutations Impair mtDNA Replication and Cause Adult-Onset Mitochondrial Encephalomyopathy. Am J Hum Genet, 97, 186–193.

49. Clausen, A.R., Lujan, S.A., Burkholder, A.B., Orebaugh, C.D., Williams, J.S., Clausen, M.F., Malc, E.P., Mieczkowski, P.A., Fargo, D.C., Smith, D.J. et al. (2015) Tracking replication enzymology *in vivo* by genome-wide mapping of ribonucleotide incorporation. Nat Struct Mol Biol, 22, 185–191.

50. Daigaku, Y., Keszthelyi, A., Muller, C.A., Miyabe, I., Brooks, T., Retkute, R., Hubank, M., Nieduszynski, C.A. and Carr, A.M. (2015) A global profile of replicative polymerase usage. Nat Struct Mol Biol, 22, 192–198.

51. Ding, J., Taylor, M.S., Jackson, A.P. and Reijns, M.A.M. (2015) Genome-wide mapping of embedded ribonucleotides and other noncanonical nucleotides using emRiboSeq and EndoSeq. Nat Protoc, 10, 1433–1444.

52. Koh, K.D., Balachander, S., Hesselberth, J.R. and Storici, F. (2015) Ribose-seq: global mapping of ribonucleotides embedded in genomic DNA. Nat Methods, 12, 251–257.

53. Reijns, M.A., Kemp, H., Ding, J., de Proce, S.M., Jackson, A.P. and Taylor, M.S. (2015) Lagging-strand replication shapes the mutational landscape of the genome. Nature, 518, 502–506.

54. Zatopek, K.M., Potapov, V., Maduzia, L.L., Alpaslan, E., Chen, L., Evans, T.C., Jr., Ong, J.L., Ettwiller, L.M. and Gardner, A.F. (2019) RADAR-seq: A RAre DAmage and Repair sequencing method for detecting DNA damage on a genome-wide scale. DNA Repair (Amst), 80, 36–44.

55. Grasso, L., Fonzino, A., Manzari, C., Leonardi, T., Picardi, E., Gissi, C., Lazzaro, F., Pesole, G. and Muzi-Falconi, M. (2024) Detection of ribonucleotides embedded in DNA by Nanopore sequencing. Commun Biol, 7, 491.

56. Wanrooij, P.H., Engqvist, M.K.M., Forslund, J.M.E., Navarrete, C., Nilsson, A.K., Sedman, J., Wanrooij, S., Clausen, A.R. and Chabes, A. (2017) Ribonucleotides incorporated by the yeast mitochondrial DNA polymerase are not repaired. Proc Natl Acad Sci U S A, 114, 12466–12471.

57. Balachander, S., Gombolay, A.L., Yang, T., Xu, P., Newnam, G., Keskin, H., El-Sayed, W.M.M., Bryksin, A.V., Tao, S., Bowen, N.E. et al. (2020) Ribonucleotide incorporation in yeast genomic DNA shows preference for cytosine and guanosine preceded by deoxyadenosine. Nature Communications, 11, 2447.

58. El-Sayed, W.M.M., Gombolay, A.L., Xu, P., Yang, T., Jeon, Y., Balachander, S., Newnam, G., Tao, S., Bowen, N.E., Bruna, T. et al. (2021) Disproportionate presence of adenosine in mitochondrial and chloroplast DNA of *Chlamydomonas reinhardtii*. iScience, 24, 102005.

59. Kundnani, D.L., Yang, T., Gombolay, A.L., Mukherjee, K., Newnam, G., Meers, C., Verma, I., Chhatlani, K., Mehta, Z.H., Mouawad, C. et al. (2024) Distinct features of ribonucleotides within genomic DNA in Aicardi-Goutieres syndrome ortholog mutants of *Saccharomyces cerevisiae*. iScience, 27, 110012.

60. Gombolay, A.L., Vannberg, F.O. and Storici, F. (2019) Ribose-Map: a bioinformatics toolkit to map ribonucleotides embedded in genomic DNA. Nucleic Acids Research, 47, e5–e5.

61. Mullakhanbhai, M.F. and Larsen, H. (1975) *Halobacterium volcanii* spec. nov., a Dead Sea halobacterium with a moderate salt requirement. Arch Microbiol, 104, 207–214.

62. Marteinsson, V.T., Birrien, J.L., Reysenbach, A.L., Vernet, M., Marie, D., Gambacorta, A., Messner, P., Sleytr, U.B. and Prieur, D. (1999) *Thermococcus barophilus* sp. nov., a new barophilic and hyperthermophilic archaeon isolated under high hydrostatic pressure from a deep-sea hydrothermal vent. Int J Syst Bacteriol, 49 Pt 2, 351–359.

63. Allers, T., Ngo, H.P., Mevarech, M. and Lloyd, R.G. (2004) Development of additional selectable markers for the halophilic archaeon *Haloferax volcanii* based on the leuB and trpA genes. Appl Environ Microbiol, 70, 943–953.

64. Birien, T., Thiel, A., Henneke, G., Flament, D., Moalic, Y. and Jebbar, M. (2018) Development of an Effective 6-Methylpurine Counterselection Marker for Genetic Manipulation in *Thermococcus barophilus*. Genes (Basel), 9, 77.

65. Meslet-Cladiere, L., Norais, C., Kuhn, J., Briffotaux, J., Sloostra, J.W., Ferrari, E., Hubscher, U., Flament, D. and Myllykallio, H. (2007) A novel proteomic approach identifies new interaction partners for proliferating cell nuclear antigen. J Mol Biol, 372, 1137–1148.

66. Godfroy, A., Raven, N.D. and Sharp, R.J. (2000) Physiology and continuous culture of the hyperthermophilic deep-sea vent archaeon *Pyrococcus abyssi* ST549. FEMS Microbiol Lett, 186, 127–132.

67. Hofer, A., Ekanem, J.T. and Thelander, L. (1998) Allosteric regulation of *Trypanosoma brucei* ribonucleotide reductase studied *in vitro* and *in vivo*. J Biol Chem, 273, 34098–34104.

68. Ranjbarian, F., Sharma, S., Falappa, G., Taruschio, W., Chabes, A. and Hofer, A. (2022) Isocratic HPLC analysis for the simultaneous determination of dNTPs, rNTPs and ADP in biological samples. Nucleic Acids Res, 50, e18.

69. Tang, M.S., Bohr, V.A., Zhang, X.S., Pierce, J. and Hanawalt, P.C. (1989) Quantification of aminofluorene adduct formation and repair in defined DNA sequences in mammalian cells using the UVRABC nuclease. J Biol Chem, 264, 14455–14462.

70. Xu, P. and Storici, F. (2021) RESCOT: Restriction Enzyme Set and Combination Optimization Tools for rNMP Capture Techniques. Theor Comput Sci, 894, 203–213.

71. Li, H., Handsaker, B., Wysoker, A., Fennell, T., Ruan, J., Homer, N., Marth, G., Abecasis, G., Durbin, R. and Genome Project Data Processing, S. (2009) The Sequence Alignment/Map format and SAMtools. Bioinformatics, 25, 2078–2079.

72. Hawkins, M., Malla, S., Blythe, M.J., Nieduszynski, C.A. and Allers, T. (2013) Accelerated growth in the absence of DNA replication origins. Nature, 503, 544–547.

73. Mc Teer, L., Moalic, Y., Cueff-Gauchard, V., Catchpole, R., Hogrel, G., Lu, Y., Laurent, S., Hemon, M., Aubé, J., Leroy, E., et al. (2024) Cooperation between two modes for DNA replication initiation in the archaeon *Thermococcus barophilus*. mBio, 15, e03200–03223.

74. Varik, V., Oliveira, S.R.A., Hauryliuk, V. and Tenson, T. (2017) HPLC-based quantification of bacterial housekeeping nucleotides and alarmone messengers ppGpp and pppGpp. Sci Rep, 7, 11022.

75. Buckstein, M.H., He, J. and Rubin, H. (2008) Characterization of nucleotide pools as a function of physiological state in *Escherichia coli*. J Bacteriol, 190, 718–726.

76. Liew, L.P., Lim, Z.Y., Cohen, M., Kong, Z., Marjavaara, L., Chabes, A. and Bell, S.D. (2016) Hydroxyurea-Mediated Cytotoxicity Without Inhibition of Ribonucleotide Reductase. Cell Rep, 17, 1657–1670.

77. Ohtani, N., Haruki, M., Morikawa, M., Crouch, R.J., Itaya, M. and Kanaya, S. (1999) Identification of the genes encoding Mn2+-dependent RNase HII and Mg2+-dependent RNase HIII from *Bacillus subtilis*: classification of RNases H into three families. Biochemistry, 38, 605–618.

78. Marinov, G.K., Bagdatli, S.T., Wu, T., He, C., Kundaje, A. and Greenleaf, W.J. (2023) The chromatin landscape of the euryarchaeon *Haloferax volcanii*. Genome Biol, 24, 253.

79. Sauguet, L., Raia, P., Henneke, G. and Delarue, M. (2016) Shared active site architecture between archaeal PolD and multi-subunit RNA polymerases revealed by X-ray crystallography. Nat Commun, 7, 12227.

80. Le Breton, M., Henneke, G., Norais, C., Flament, D., Myllykallio, H., Querellou, J. and Raffin, J.P. (2007) The heterodimeric primase from the euryarchaeon *Pyrococcus abyssi*: a multifunctional enzyme for initiation and repair? J Mol Biol, 374, 1172–1185.

81. Martinez-Carranza, M., Vialle, L., Madru, C., Cordier, F., Tekpinar, A.D., Haouz, A., Legrand, P., Le Meur, R.A., England, P., Dulermo, R., et al. (2024) Communication between DNA polymerases and Replication Protein A within the archaeal replisome. Nat Commun, 15, 10926.

82. Malfatti, M.C., Henneke, G., Balachander, S., Koh, K.D., Newnam, G., Uehara, R., Crouch, R.J., Storici, F. and Tell, G. (2019) Unlike the *Escherichia coli* counterpart, archaeal RNase HII cannot process ribose monophosphate abasic sites and oxidized ribonucleotides embedded in DNA. J Biol Chem, 294, 13061–13072.

83. Grant, J.R., Enns, E., Marinier, E., Mandal, A., Herman, E.K., Chen, C.Y., Graham, M., Van Domselaar, G. and Stothard, P. (2023) Proksee: in-depth characterization and visualization of bacterial genomes. Nucleic Acids Res, 51, W484–W492.

84. Ignatov, K.B., Blagodatskikh, K.A., Shcherbo, D.S., Kramarova, T.V., Monakhova, Y.A. and Kramarov, V.M. (2019) Fragmentation Through Polymerization (FTP): A new method to fragment DNA for next-generation sequencing. PLoS One, 14, e0210374.

