## Supplementary material for "Genome-wide ribonucleotide detection in *Archaea*": Tables+Supplemental_Tables

### SUPPLEMENTARY DATA

**Supplementary Figure S1.** Distribution of RE cutting sites generated by *in silico* digestion across the chromosomes.
**Supplementary Figure S2.** Distribution of ribonucleotide sites on Hvo Plasmids.
**Supplementary Figure S3.** Distribution of DNA base sequences around rNMP incorporation sites in pHV1 in all libraries of Hvo strains.
**Supplementary Figure S4.** Distribution of DNA base sequences around rNMP incorporation sites in pHV3 in all libraries of Hvo strains.
**Supplementary Figure S5.** Distribution of DNA base sequences around rNMP incorporation sites in pHV4 in all libraries of Hvo strains.
**Supplementary Figure S6.** Distribution of DNA base sequences at rNMP incorporation sites.
**Supplementary Figure S7.** Structural comparison of type 2 RNases H. (A) Domain architecture type 2 RNase H proteins.
**Supplementary Figure S8.** Structural comparison of type 1 RNases H. (A) Domain architecture type 1 RNase H proteins.
**Supplementary Figure S9.** Comparison of type 2 RNases H.
**Supplementary Figure S10.** Comparison of type 1 RNases H.
**Supplementary Table S1**. Genomic architecture of the archaeal biological models used in this study.
**Supplementary Table S2**. ribose-seq datasets generated in this study.
**Supplementary Table S3.** Strand incorporation values of ribose-seq (R2 datatsets).
**Supplementary Table S4.** Strand incorporation values of ribose-seq (R1 datasets).
**SupplementaryTable S5.** Rate of rNMP incorporation (%) for the R1 datasets.

### SUPPLEMENTARY EXPERIMENTAL PROCEDURES

### SUPPLEMENTARY REFERENCES

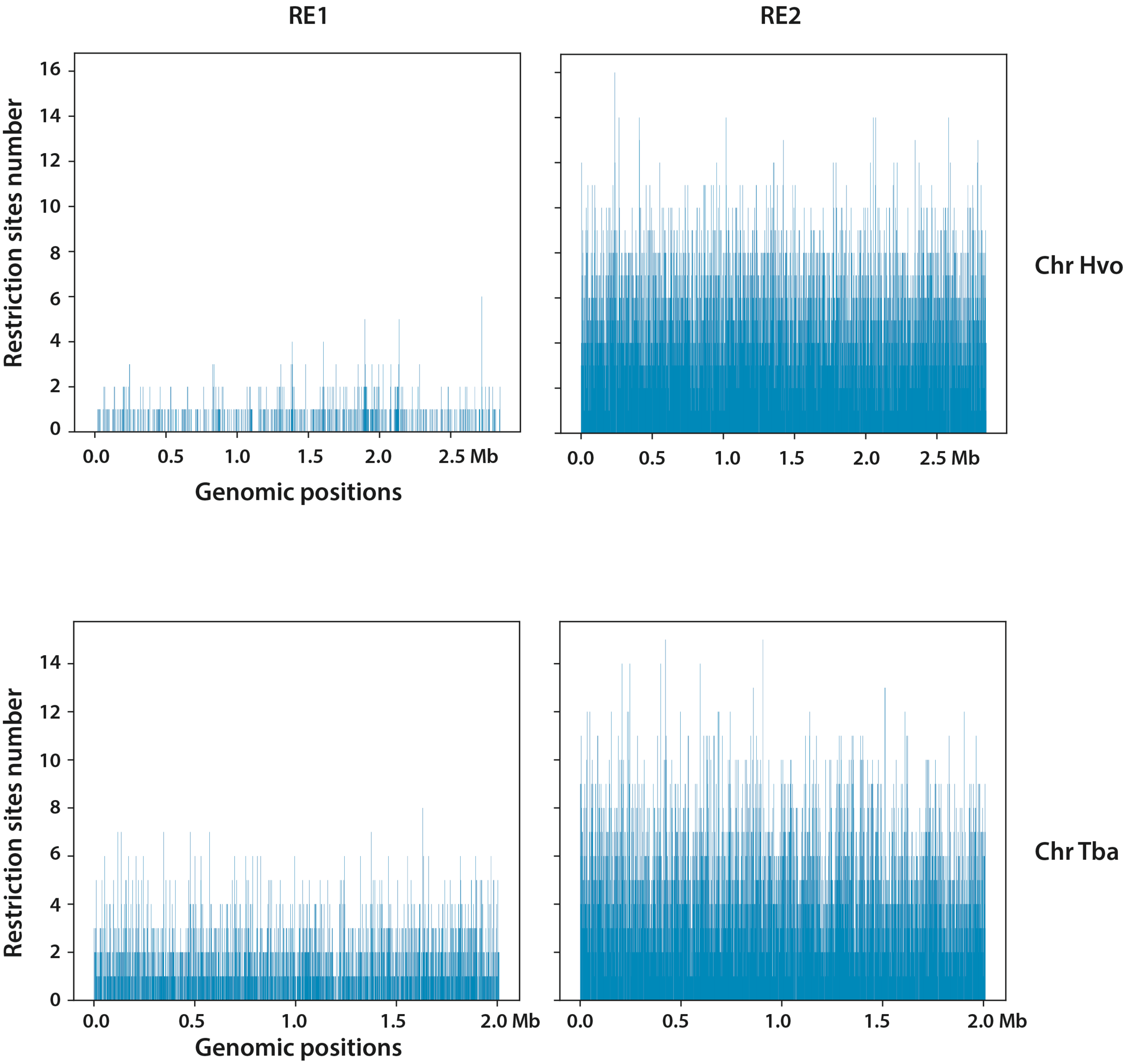
RE1
RE2
Chr Hvo
Chr Tba
Restriction sites number
Genomic positions
0.0
0.5
1.0
1.5
2.0
2.5 Mb
2.0 Mb
0
2
4
6
8
10
12
14
16

**Figure S1. Distribution of RE cutting sites generated by *in silico* digestion across the chromosomes**. The number and position of cutting sites according to the cocktails of restriction enzymes used in RE1 (*DraI*, *EcoRV,* and *SspI*) or RE2 (*MspA1I*, *HpyCH4V*) are determined for each chromosome species (Hvo and Tba). Histogram plots of RE cutting site positions within 2 kb-sized bins are shown.

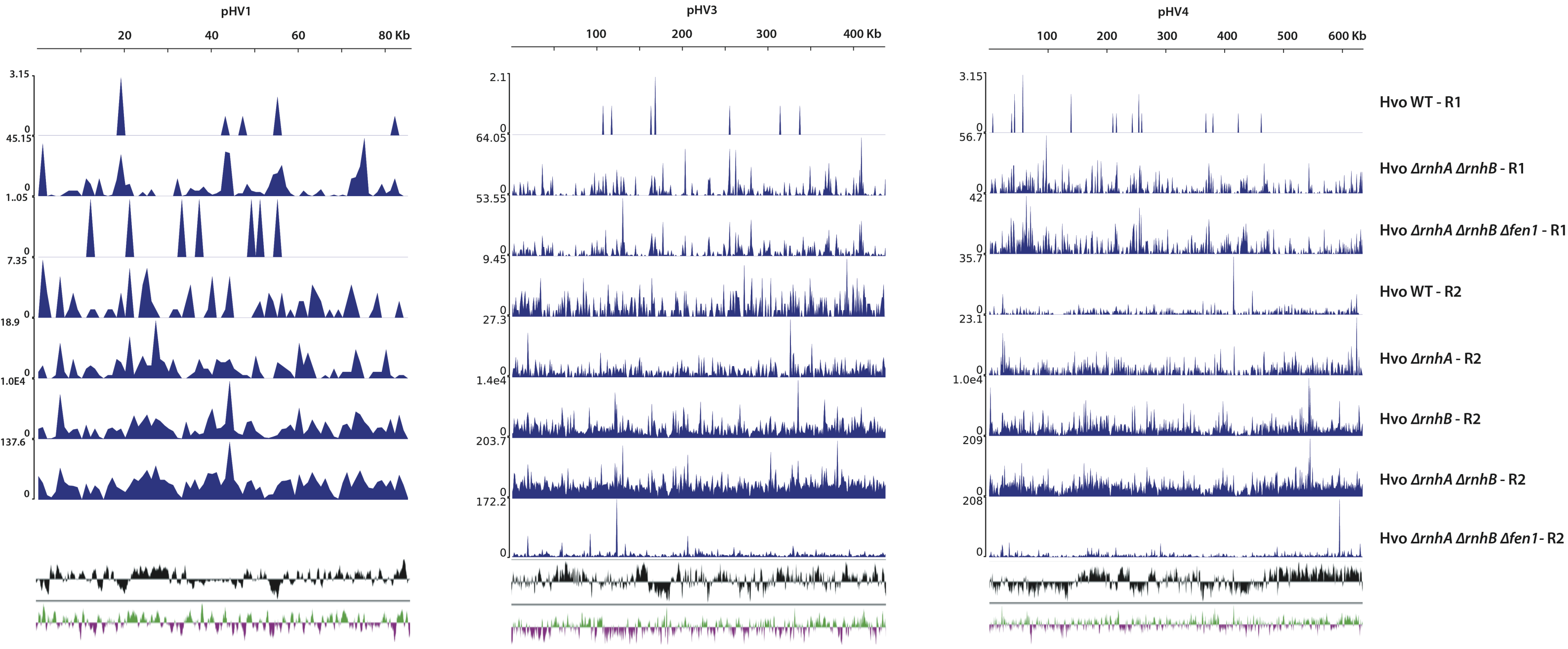
pHV1
20
40
60
80 Kb
pHV3
100
200
300
400 Kb
pHV4
100
200
300
400
500
600 Kb
Hvo WT - R1
Hvo ΔrnhA ΔrnhB - R1
Hvo ΔrnhA ΔrnhB Δfen1 - R1
Hvo WT - R2
Hvo ΔrnhA - R2
Hvo ΔrnhB - R2
Hvo ΔrnhA ΔrnhB - R2
Hvo ΔrnhA ΔrnhB Δfen1- R2

**Figure S2**. **Distribution of ribonucleotide sites on Hvo Plasmids.** Locations and occurrences of rNMPs in the genetic elements pHV1, pHV3 and pHV4 of *H. volcanii* strains (Hvo) from two ribose-seq rounds (R1 and R2). rNMP distributions across plasmids of wild-type and mutant Hvo strains are shown. Occurrence of rNMPs is plotted in every 1 kb-sized plasmid sequences.

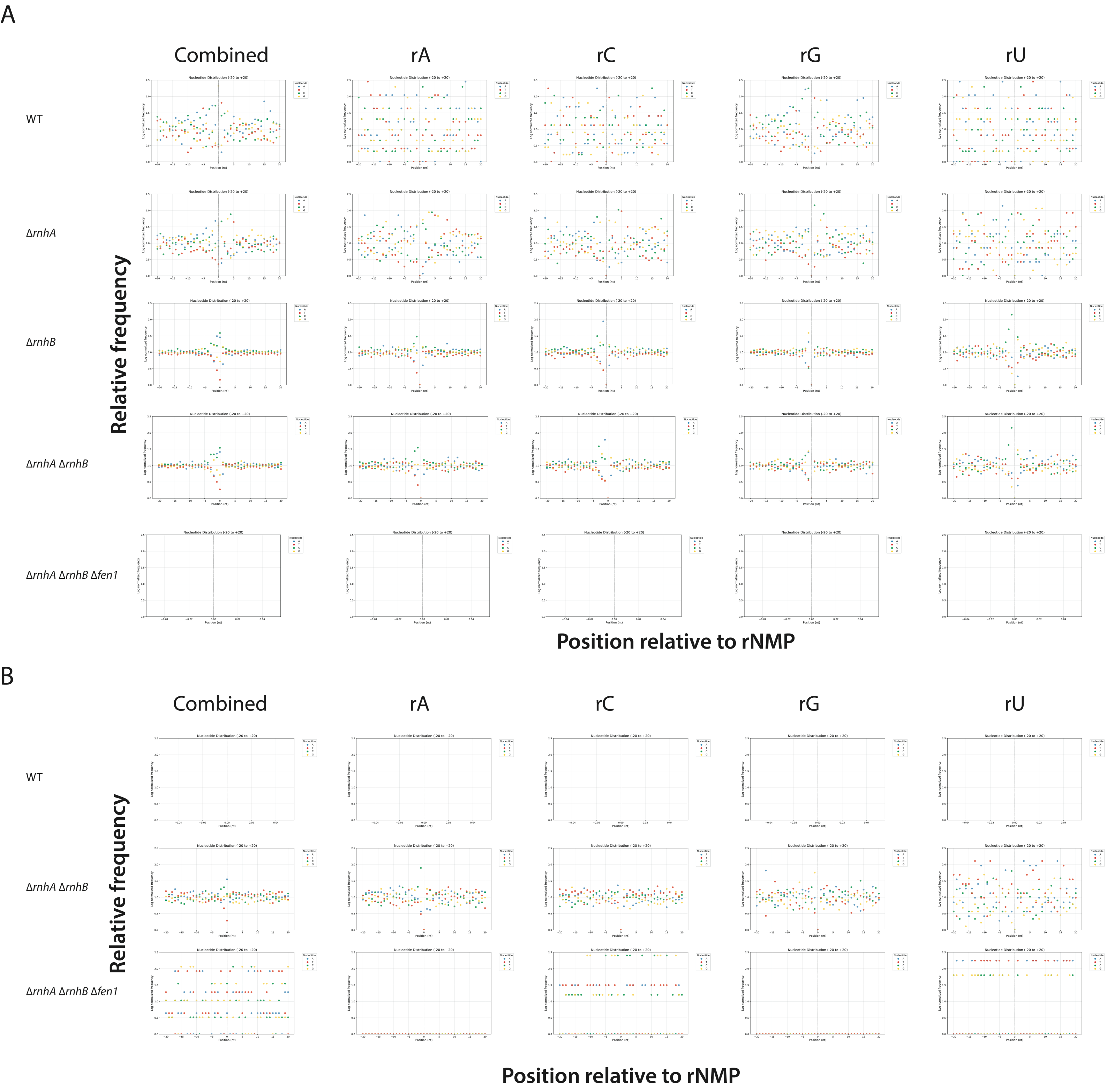

A
Combined
rA
rC
rG
rU
WT
ΔrnhA
ΔrnhB
ΔrnhA ΔrnhB
ΔrnhA ΔrnhB Δfen1
Relative frequency
Position relative to rNMP
B
Combined
rA
rC
rG
rU
WT
ΔrnhA ΔrnhB
ΔrnhA ΔrnhB Δfen1
Relative frequency
Position relative to rNMP

**Figure S3. Distribution of DNA base sequences around rNMP incorporation sites in pHV1 in all libraries of Hvo strains.** R2 datatsets **(A)** and R1 datasets **(B)** are shown. Base-distribution sequences encountered at the genomic coordinate of the rNMP incorporation site are computed for all four nucleotides (“combined”) and separately for each ribonucleotide (rA, rC, rG and rU). The forward and reverse strands are not differentiated.

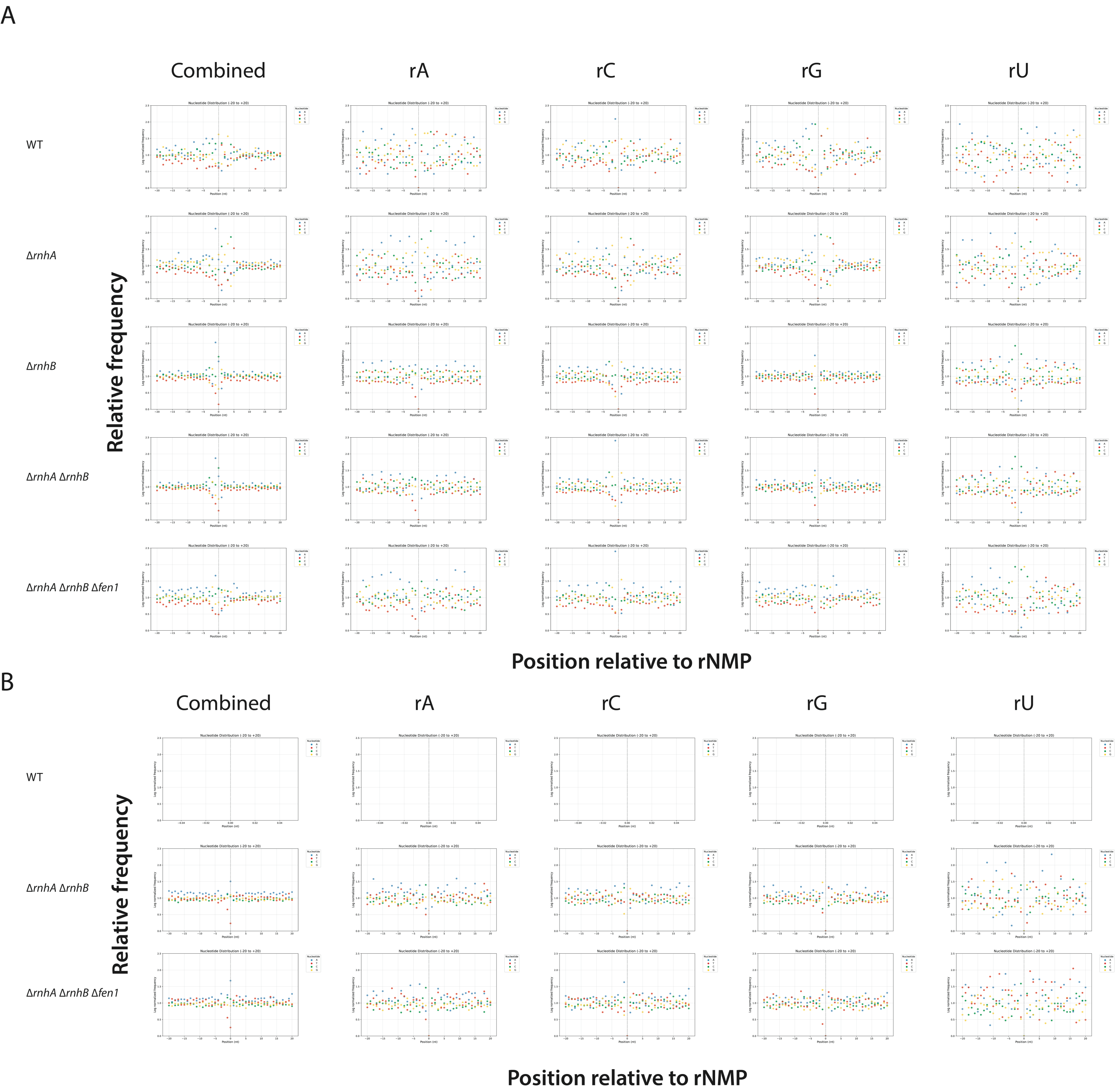
A
Combined
rA
rC
rG
rU
WT
ΔrnhA
ΔrnhB
ΔrnhA ΔrnhB
ΔrnhA ΔrnhB Δfen1
Relative frequency
Nucleotide Distribution (-20 to +20)
Position relative to rNMP
B
Combined
rA
rC
rG
rU
WT
ΔrnhA ΔrnhB
ΔrnhA ΔrnhB Δfen1
Relative frequency
Position relative to rNMP

**Figure S4. Distribution of DNA base sequences around rNMP incorporation sites in pHV3 in all libraries of Hvo strains.** R2 datatsets **(A)** and R1 datasets **(B)** are shown. Base-distribution sequences encountered at the genomic coordinate of the rNMP incorporation site are computed for all four nucleotides ("combined") and separately for each ribonucleotide (rA, rC, rG and rU). The forward and reverse strands are not differentiated.

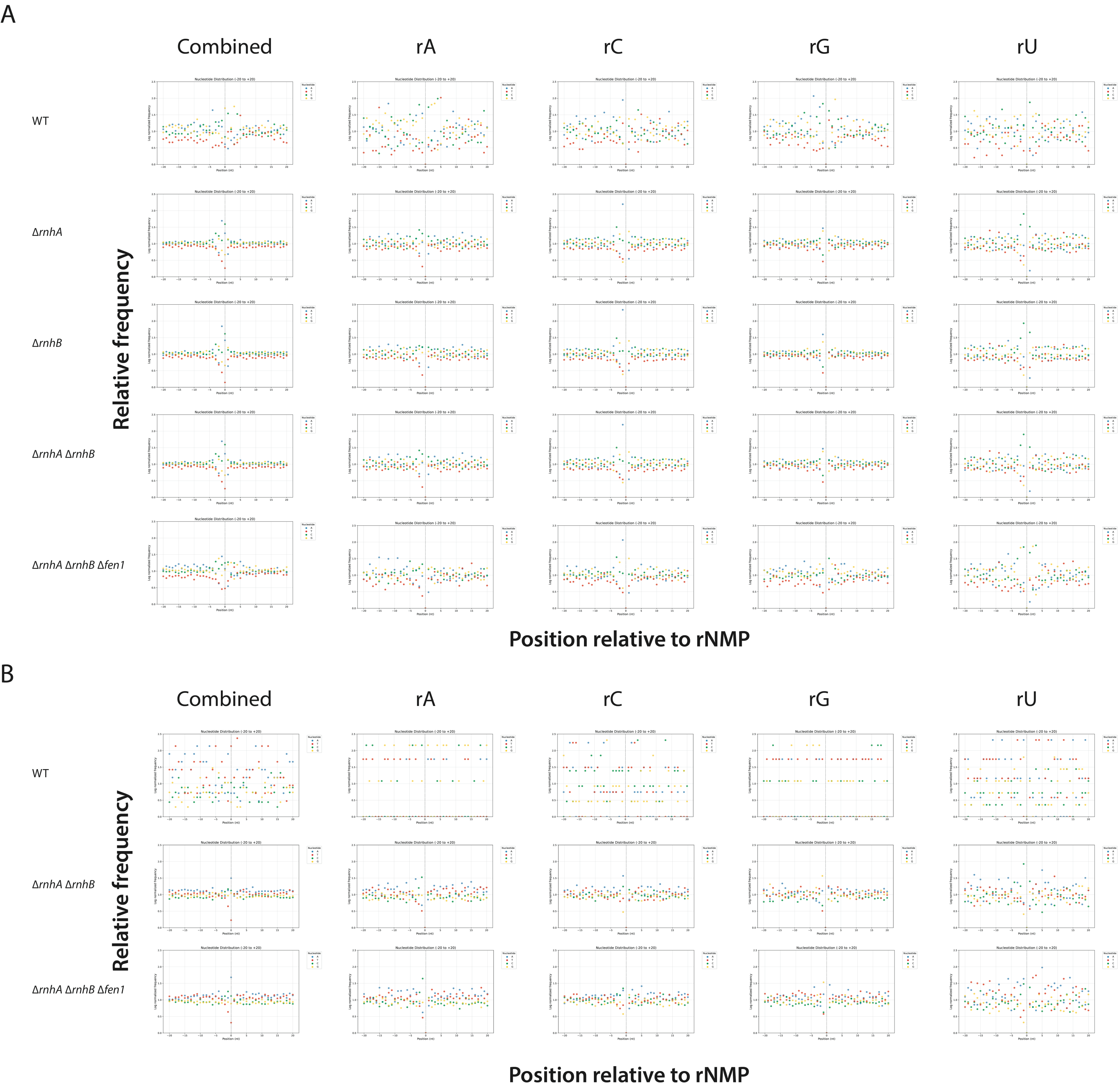

A
Combined
rA
rC
rG
rU
WT
ΔrnhA
ΔrnhB
ΔrnhA ΔrnhB
ΔrnhA ΔrnhB Δfen1
Relative frequency
Position relative to rNMP
B
Combined
rA
rC
rG
rU
WT
ΔrnhA ΔrnhB
ΔrnhA ΔrnhB Δfen1
Relative frequency
Position relative to rNMP

**Figure S5. Distribution of DNA base sequences around rNMP incorporation sites in pHV4 in all libraries of Hvo strains.** R2 datatsets **(A)** and R1 datasets **(B)** are shown. Base-distribution sequences encountered at the genomic coordinate of the rNMP incorporation site are computed for all four nucleotides ("combined") and separately for each ribonucleotide (rA, rC, rG and rU). The forward and reverse strands are not differentiated.

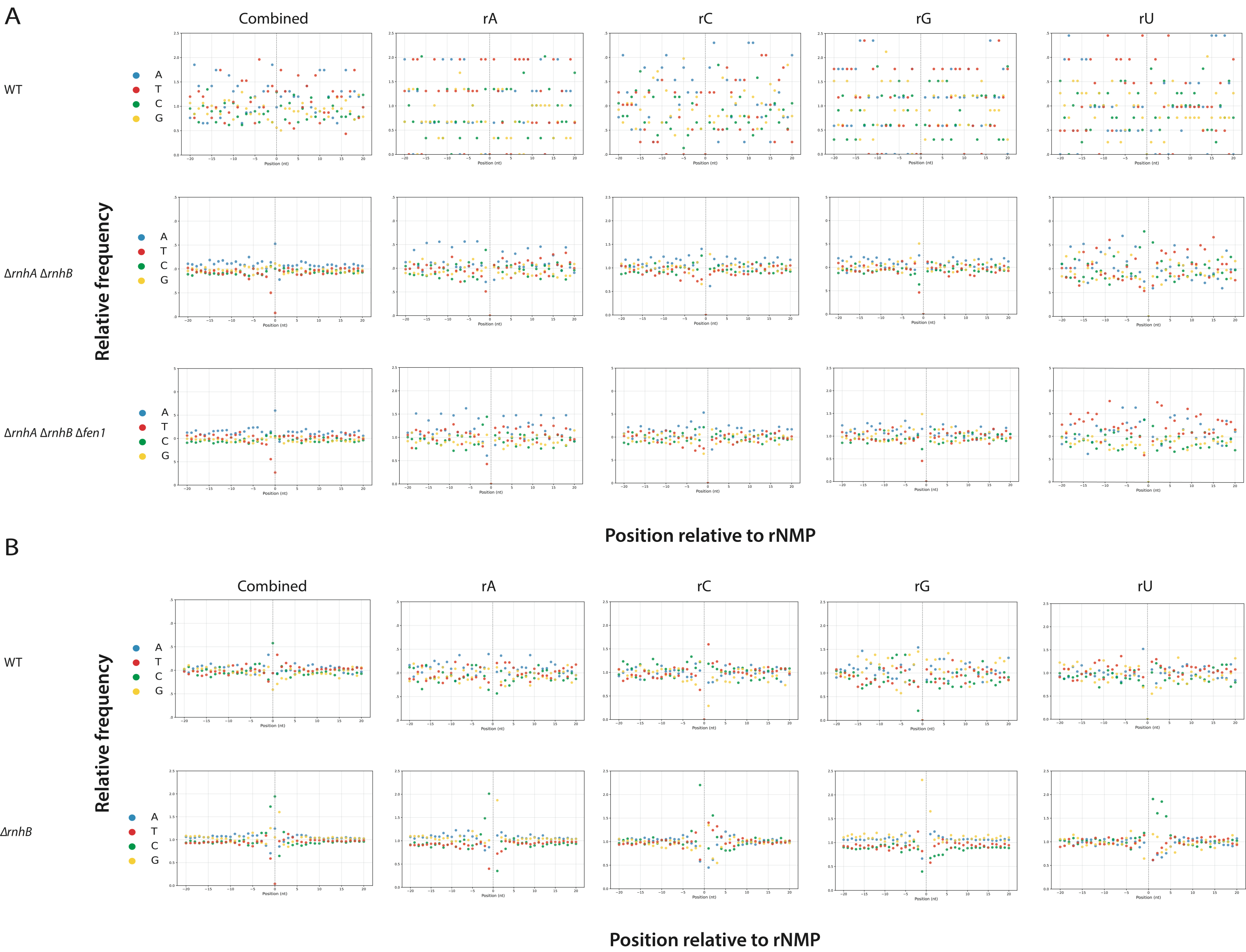

A
Combined
rA
rC
rG
rU
WT
ΔrnhA ΔrnhB
ΔrnhA ΔrnhB Δfen1
A
T
C
G
Relative frequency
Position (nt)
Position relative to rNMP
B
Combined
rA
rC
rG
rU
WT
ΔrnhB
Relative frequency
Position relative to rNMP

**Figure S6. Distribution of DNA base sequences at rNMP incorporation sites. (A)** Chromosome of Hvo strains. **(B)** Chromosome of Tba strains. For each strain, the base distribution encountered at the genomic coordinate of the rNMP incorporation site is computed for all four rNMP ("combined") and separately for each rNMP (rA, rC, rG and rU). Position 0 on the x-axis represents the site of rNMP incorporation, − and + positions represent upstream and downstream dNMPs, respectively. The y-axis shows the frequency of each base found in the genomic context surrounding the rNMP incorporation sites normalized to the frequency of the corresponding base in the reference genome. R1 datasets are shown.

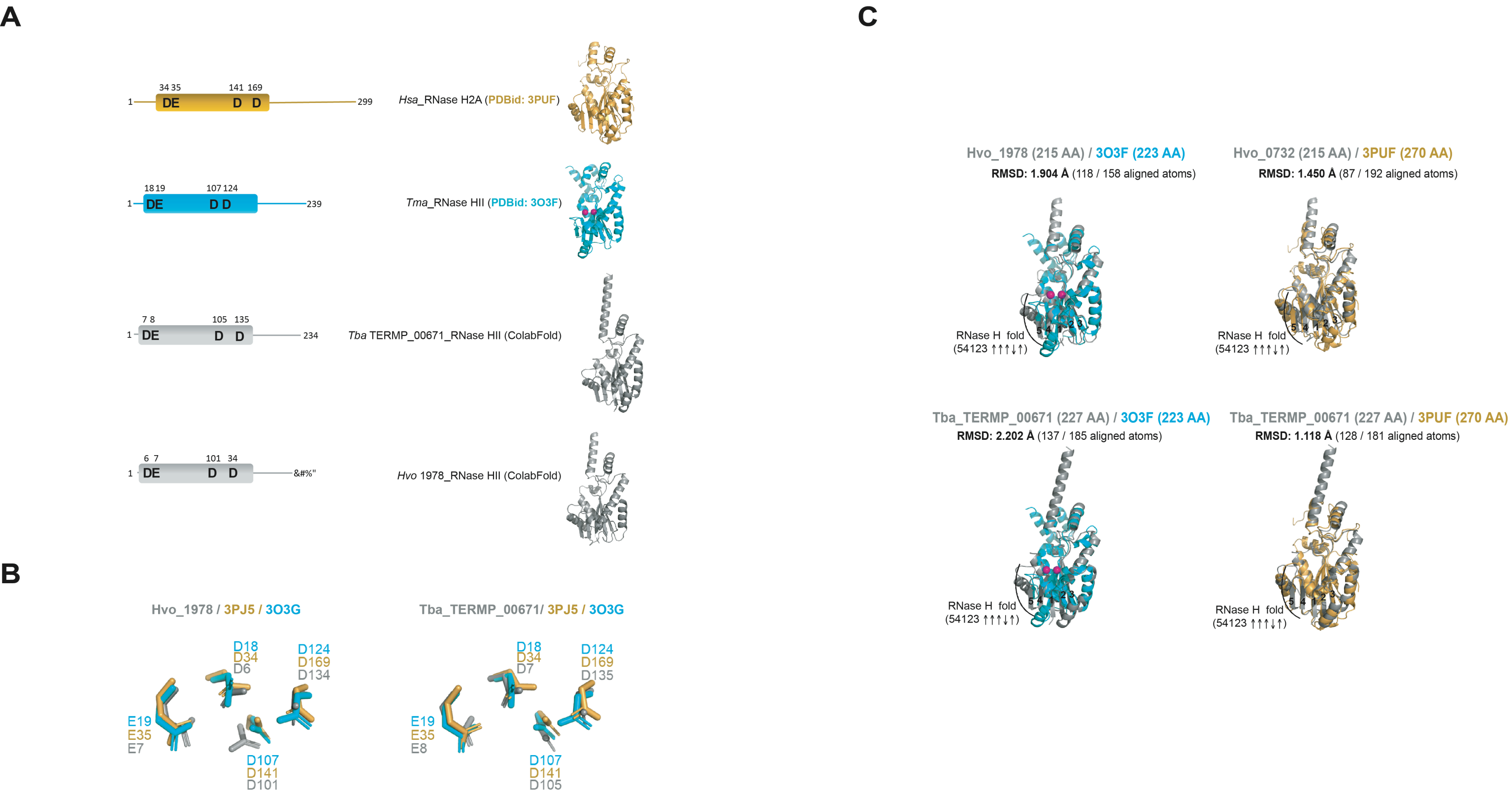
A
B
C
34 35
141 169
DE
D D
1
299
Hsa_RNase H2A (PDBid: 3PUF)
18 19
107 124
239
Tma_RNase HII (PDBid: 3O3F)
7 8
105
135
234
Tba TERMP_00671_RNase HII (ColabFold)
6 7
101
34
&#%"
Hvo 1978_RNase HII (ColabFold)
Hvo_1978 / 3PJ5 / 3O3G
Tba_TERMP_00671/ 3PJ5 / 3O3G
D18
D34
D6
D124
D169
D134
E19
E35
E7
D107
D141
D101
D7
D135
E8
D105
Hvo_1978 (215 AA) / 3O3F (223 AA)
RMSD: 1.904 Å (118 / 158 aligned atoms)
Hvo_0732 (215 AA) / 3PUF (270 AA)
RMSD: 1.450 Å (87 / 192 aligned atoms)
Tba_TERMP_00671 (227 AA) / 3O3F (223 AA)
RMSD: 2.202 Å (137 / 185 aligned atoms)
Tba_TERMP_00671 (227 AA) / 3PUF (270 AA)
RMSD: 1.118 Å (128 / 181 aligned atoms)
RNase H fold
(54123 ↑↑↑↓↑)

**Figure S7. Structural comparison of type 2 RNases H. (A) Domain architecture type 2 RNase H proteins**. The conserved structural core of the RNase H domain is aligned between all proteins and the four active site residues are indicated. **(B) Overlay of the active site of type 2 RNase H proteins**. On the left, a zoom-in view of the active site of Hvo_1978 RNase HII and the two experimentally solved *Tma*_RNase HII (PDBid: 3O3G) and *Hsa*_RNase H1 (PDBid: 3PJ5) structures. Superposition of the active sites with respect to Hvo_1978 RNase HII predicted structure. On the right, a close up view of Tba_TERMP_00671 RNase HII and the two experimentally solved *Tma*_RNase HII (PDBid: 3O3G) and *Hsa*_RNase H2A (PDBid: 3PJ5) structures. Superposition of the active sites with respect to Tba_TERMP_00671 RNase HII predicted structure. **(C) Superimposed 3-D structure of type 2 RNase H proteins.** On the left, Hvo_1978 RNase HII predicted structure (215 amino acids) is represented in grey and the experimentally solved *Tma*_RNase HII structure (223 amino acids, PDBid: 3O3F) is in cyan. The conserved RNase H fold consisting of a five-stranded β sheet is shown. RMSD value (in Å) of C-alpha atoms of superimposed structures is shown. On the right, Hvo_1978 RNase HII predicted structure (215 amino acids) is represented in grey and the experimentally solved *Hsa*_RNase H2A structure (270 amino acids, PDBid: 3PUF, chain A) is in bright-orange. The conserved RNase H fold consisting of a five-stranded β sheet is indicated. RMSD value (in Å) of C-alpha atoms of superimposed structures is reported. On the middle left, Tba_TERMP_00671 RNase HII predicted structure (227 amino acids) is represented in grey and the experimentally solved *Tma*_RNase HII structure (223 amino acids, PDBid: 3O3F) is in cyan. The conserved RNase H fold consisting of a five-stranded β sheet is shown. RMSD value (in Å) of C-alpha atoms of superimposed structures is shown. On the middle right, Tba_TERMP_00671 RNase HII predicted structure (227 amino acids) is represented in grey and the experimentally solved *Hsa*_RNase H1 structure (270 amino acids, PDBid: 3PUF, chain A) is in bright-orange. The conserved RNase H fold consisting of a five-stranded β sheet is indicated. RMSD value (in Å) of C-alpha atoms of superimposed structures is reported.

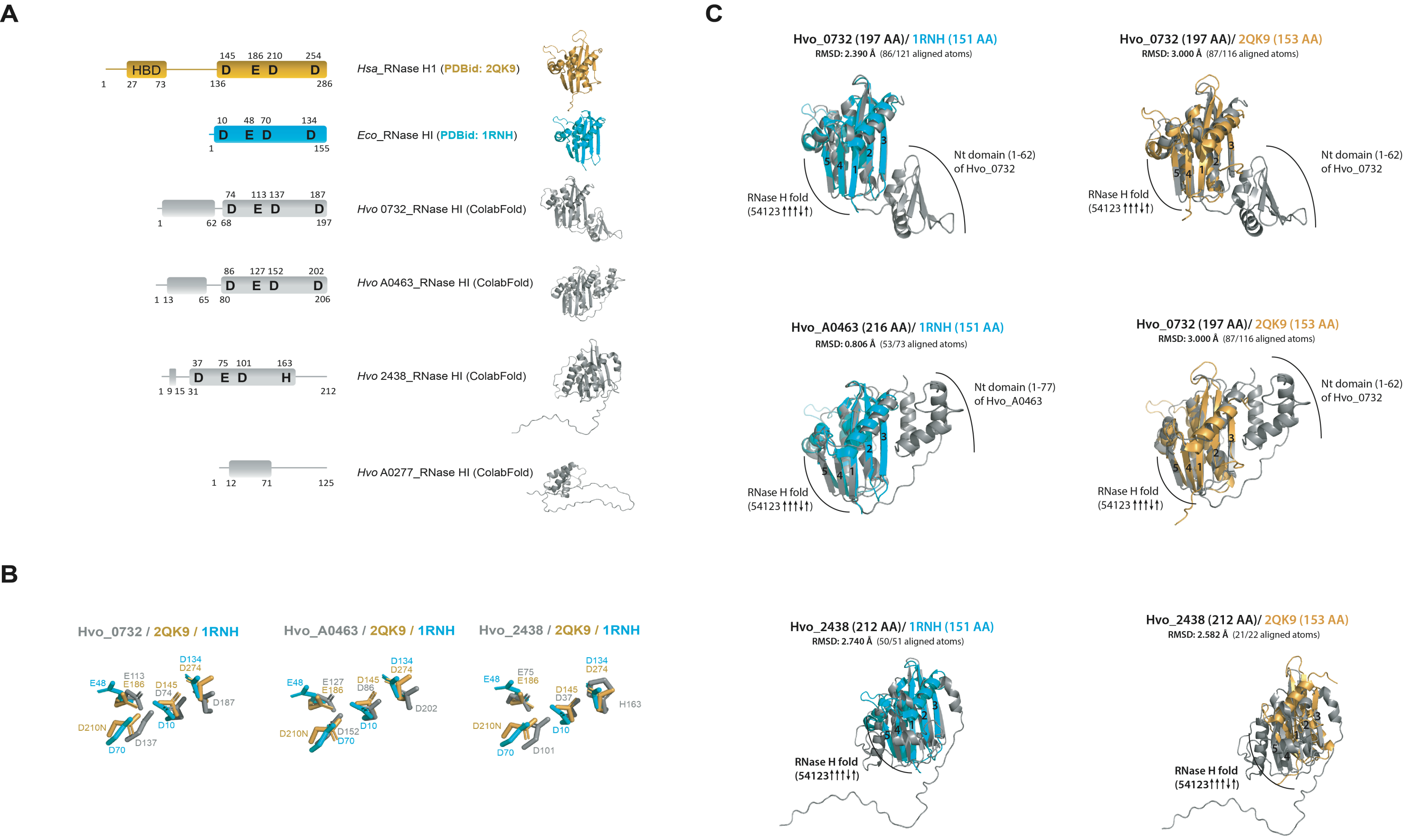

A
HBD
145 186 210 254
D E D D
1 27 73 136 286
Hsa_RNase H1 (PDBid: 2QK9)
10 48 70 134
D E D D
1 155
Eco_RNase HI (PDBid: 1RNH)
74 113 137 187
D E D D
1 62 68 197
Hvo 0732_RNase HI (ColabFold)
86 127 152 202
D E D D
1 13 65 80 206
Hvo A0463_RNase HI (ColabFold)
37 75 101 163
D E D H
1 9 15 31 212
Hvo 2438_RNase HI (ColabFold)
1 12 71 125
Hvo A0277_RNase HI (ColabFold)
B
Hvo_0732 / 2QK9 / 1RNH
E48 E113 E186 D134 D274 D145 D74 D187 D210N D10 D70 D137
Hvo_A0463 / 2QK9 / 1RNH
E48 E127 E186 D134 D274 D145 D86 D202 D210N D152 D10 D70
Hvo_2438 / 2QK9 / 1RNH
E48 E75 E186 D134 D274 D145 D37 H163 D210N D10 D70 D101
C
Hvo_0732 (197 AA)/ 1RNH (151 AA)
RMSD: 2.390 Å (86/121 aligned atoms)
Nt domain (1-62) of Hvo_0732
RNase H fold (54123↑↑↑↓↑)
Hvo_0732 (197 AA)/ 2QK9 (153 AA)
RMSD: 3.000 Å (87/116 aligned atoms)
Nt domain (1-62) of Hvo_0732
RNase H fold (54123↑↑↑↓↑)
Hvo_A0463 (216 AA)/ 1RNH (151 AA)
RMSD: 0.806 Å (53/73 aligned atoms)
Nt domain (1-77) of Hvo_A0463
RNase H fold (54123↑↑↑↓↑)
Hvo_0732 (197 AA)/ 2QK9 (153 AA)
RMSD: 3.000 Å (87/116 aligned atoms)
Nt domain (1-62) of Hvo_0732
RNase H fold (54123↑↑↑↓↑)
Hvo_2438 (212 AA)/ 1RNH (151 AA)
RMSD: 2.740 Å (50/51 aligned atoms)
RNase H fold (54123↑↑↑↓↑)
Hvo_2438 (212 AA)/ 2QK9 (153 AA)
RMSD: 2.582 Å (21/22 aligned atoms)
RNase H fold (54123↑↑↑↓↑)

**Figure S8. Structural comparison of type 1 RNases H. (A) Domain architecture type 1 RNase H proteins**. The conserved structural core of the RNase H domain is aligned between all proteins and the four active site residues are indicated. For *Hsa*_RNase H1, the Hybrid Binding Domain (HBD) is represented. When relevant, secondary structural elements are schematically depicted. **(B) Overlay of the active site of type 1 RNase H proteins**. To the left, a zoom-in view of the active site of Hvo_0732 RNase HI and the two experimentally solved *Eco*_RNase HI (PDBid: 1RNH) and *Hsa*_RNase H1 (PDBid: 2QK9) structures. To the middle, a close up view of Hvo_A0463 RNases HI and the two experimentally solved *Eco*_RNase HI (PDBid: 1RNH) and *Hsa*_RNase H1 (PDBid: 2QK9) structures. To the right, a close up view of Hvo_2468 RNase HI and the two experimentally solved *Eco*_RNase HI (PDBid: 1RNH) and *Hsa*_RNase H1 (PDBid: 2QK9) structures. 1RNH was used as PDB reference for the superposition of the active sites. **(C) Superimposed 3-D structure of type 1 RNase H proteins.** On the left, Hvo_0732 RNase HI predicted structure (197 amino acids) is represented in grey and the experimentally solved *Eco*_RNase HI structure (151 amino acids, PDBid: 1RNH) is in cyan. The conserved RNase H fold consisting of a five-stranded β sheet is shown and the N-terminal domain of Hvo_0732 RNase HI is indicated. RMSD (root-mean-square deviation – in Å) value of C-alpha atoms of superimposed structures is shown. On the right, Hvo_0732 RNase HI predicted structure (197 amino acids) is represented in grey and the experimentally solved *Hsa*_RNase H1 structure (153 amino acids, PDBid: 2QK9) is in bright-orange. The conserved RNase H fold consisting of a five-stranded β sheet is shown and the N-terminal domain of Hvo_0732 RNase HI is indicated. RMSD value (in Å) of C-alpha atoms of superimposed structures is reported. On the middle left, Hvo_A0463 RNase HI predicted structure (216 amino acids) is represented in grey and the experimentally solved *Eco*_RNase HI structure (151 amino acids, PDBid: 1RNH) is in cyan. The conserved RNase H fold consisting of a five-stranded β sheet is shown and the N-terminal domain of Hvo_A0463 RNase HI is indicated. RMSD value (in Å) of C-alpha atoms of superimposed structures is shown. On the middle right, Hvo_A0463 RNase HI predicted structure (216 amino acids) is represented in grey and the experimentally solved *Hsa*_RNase H1 structure (153 amino acids, PDBid: 2QK9) is in bright-orange. The conserved RNase H fold consisting of a five-stranded β sheet is shown and the N-terminal domain of Hvo_A0463 RNase HI is indicated. RMSD value (in Å) of C-alpha atoms of superimposed structures is reported. On the bottom left, Hvo_2438 RNase HI predicted structure (212 amino acids) is represented in grey and the experimentally solved *Eco*_RNase HI structure (151 amino acids, PDBid: 1RNH) is in cyan. The conserved RNase H fold consisting of a five-stranded β sheet is shown. RMSD value (in Å) of C-alpha atoms of superimposed structures is shown. On the middle right, Hvo_2438 RNase HI predicted structure (212 amino acids) is represented in grey and the experimentally solved *Hsa*_RNase H1 structure (153 amino acids, PDBid: 2QK9) is in bright-orange. The conserved RNase H fold consisting of a five-stranded β sheet is shown. RMSD value (in Å) of C-alpha atoms of superimposed structures is reported.

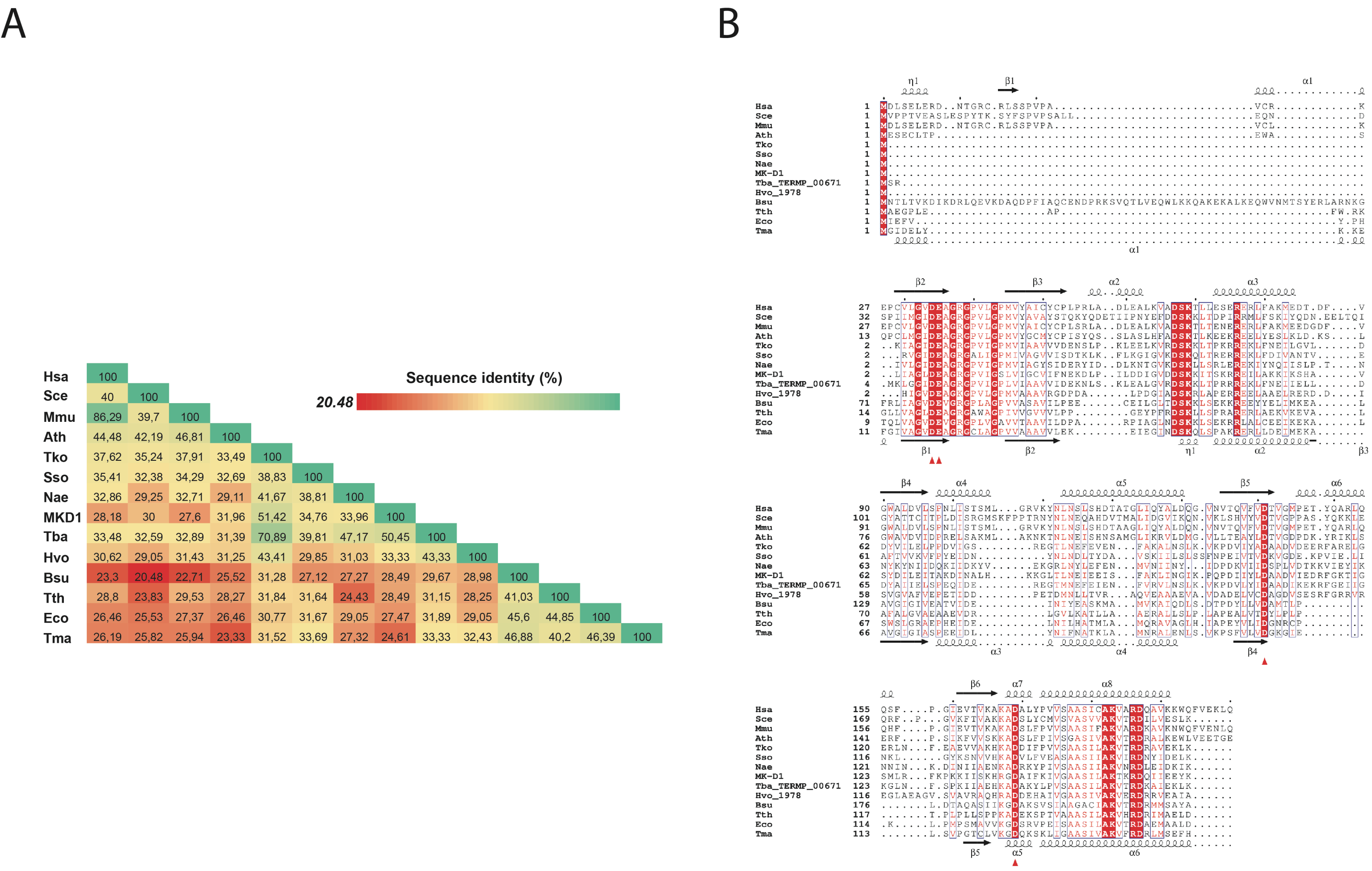
A
Sequence identity (%)
20.48
Hsa 100
Sce 40 100
Mmu 86,29 39,7 100
Ath 44,48 42,19 46,81 100
Tko 37,62 35,24 37,91 33,49 100
Sso 35,41 32,38 34,29 32,69 38,83 100
Nae 32,86 29,25 32,71 29,11 41,67 38,81 100
MKD1 28,18 30 27,6 31,96 51,42 34,76 33,96 100
Tba 33,48 32,59 32,89 31,39 70,89 39,81 47,17 50,45 100
Hvo 30,62 29,05 31,43 31,25 43,41 29,85 31,03 33,33 43,33 100
Bsu 23,3 20,48 22,71 25,52 31,28 27,12 27,27 28,49 29,67 28,98 100
Tth 28,8 23,83 29,53 28,27 31,84 31,64 24,43 28,49 31,15 28,25 41,03 100
Eco 26,46 25,53 27,37 26,46 30,77 31,67 29,05 27,47 31,89 29,05 45,6 44,85 100
Tma 26,19 25,82 25,94 23,33 31,52 33,69 27,32 24,61 33,33 32,43 46,88 40,2 46,39 100
B
Hsa
Sce
Mmu
Ath
Tko
Sso
Nae
MK-D1
Tba_TERMP_00671
Hvo_1978
Bsu
Tth
Eco
Tma

**Figure S9. Comparison of type 2 RNases H. (A)** Pairwise identity matrix of *Homo sapiens* (Hsa), *Saccharomyces cerevisiae* (Sce), *Mus musculus* (Mmu), *Arabidopsis thaliana* (Ath), *Thermococcus kodakarensis* strain KOD1 (Tko), *Saccharolobus solfataricus* strain P2 (Sso), *Nanobdella aerobiophila* (Nae), *Candidatus Prometheoarchaeum syntrophicum* strain MK-D1 (MK-D1), *Thermococcus barophilus* (Tba_TERMP_00671), *Haloferax volcanii* strain H53 (Hvo_1978), *Bacillus subtilis* (Bsu), *Thermus thermophilus* (Tth), *Escherichia coli* (Eco) and *Thermotoga maritima* (Tma). The colour represents percent identity, from lowest 20.48% (red) to highest 100% (green). **(B)** Structure-based sequence alignment is based on the three-dimensional structures of RNase H2A_Hsa (corresponding structure elements are shown at the top)(PDBid: 3PUF) and RNase HII_Tma (corresponding structure elements are shown at the bottom)(PDBid: 3O3F). Conserved active site residue and the tyrosine are marked with red arrows and blue dot, respectively. Similar residues are in red. Strictly conserved residues are highlighted in red.

A

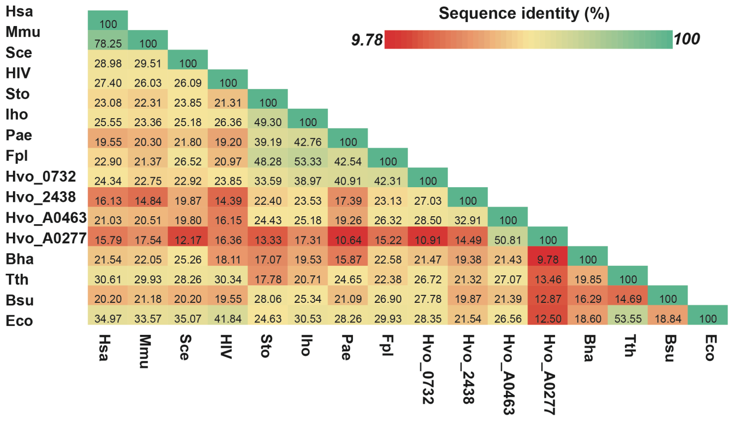
Sequence identity (%)
9.78
100
Hsa
Mmu
Sce
HIV
Sto
Iho
Pae
Fpl
Hvo_0732
Hvo_2438
Hvo_A0463
Hvo_A0277
Bha
Tth
Bsu
Eco

B

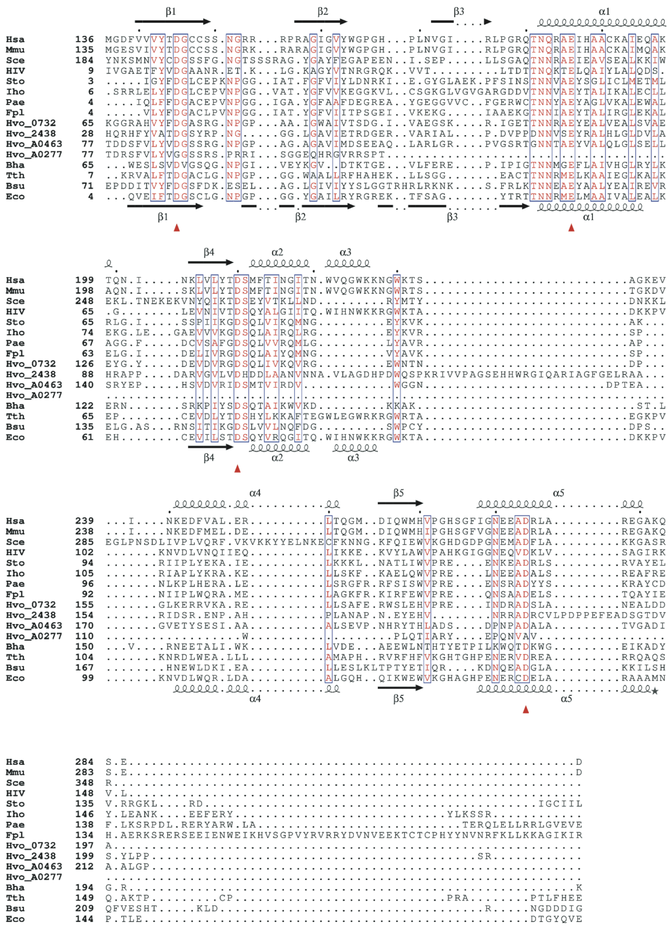
β1
β2
β3
α1
β4
α2
α3
α4
β5
α5

**Figure S10. Comparison of type 1 RNases H. (A)** Pairwise identity matrix of *Homo sapiens* (Hsa), *Mus musculus* (Mmu), *Saccharomyces cerevisiae* (Sce), HIV (Human Immunodeficiency Virus RT), *Sulfurisphaera tokodaii* (Sto), *Ignicoccus hospitalis* strain KIN4/I (Iho), *Pyrobaculum aerophilum* strain IM2 (Pae), *Ferroglobus placidus* strain DSM 10642, *Haloferax volcanii* strain H53 chromosome-encoded RNase H1_0732 (Hvo_0732), *Haloferax volcanii* strain H53 chromosome-encoded RNase H1_2438 (Hvo_2438), *Haloferax volcanii* strain H53 plasmid-encoded RNase H1_A0463 (Hvo_0463), *Haloferax volcanii* strain H53 plasmid-encoded RNase H1_A0277 (Hvo_A0277), *Bacillus halodurans* (Bha), *Thermus thermophilus* (Tth), *Bacillus subtilis* (Bsu) and *Escherichia coli* (Eco). The colour represents percent identity, from lowest 9.78% (red) to highest 100% (green). **(B)** Structure-based sequence alignment is based on the three-dimensional structures of RNase H1_Hsa (corresponding structure elements are shown at the top)(PDBid: 2QK9) and RNase HI_Eco (corresponding structure elements are shown at the bottom)(PDBid: 1RNH). Conserved active site residues are marked with red arrows. Similar residues are in red.

**Table S1. Genomic architecture of the archaeal biological models used in this study.**

| **organism** | *Haloferax volcanii* H53 | | | | *Thermococcus barophilus* MP |
|---|---|---|---|---|---|
| **Genetic element** | Chr | pHV1 | pHV3 | pHV4 | Chr |
| **NCBI Reference Sequence** | NC_013967.1 | NC_013968.1 | NC_013964.1 | NC_013966.1 | NC_014804.1 |
| **Genome size (bp)** | 2,847,757 | 85,092 | 437,906 | 635,786 | 2,010,078 |
| **GC%** | 66.7 | 56 | 66 | 62 | 42 |

**Table S2. ribose-seq datasets generated in this study.** Genomic coordinates are the number of different locations of rNMPs detected on the plasmids while occurrences are the overall rNMPs detected. (N/A stands for data missing after the ribose-map process)

| Library | Organism | Genotype | Genomic coordinates | Occurrences | Restriction Enzymes | ribose-seq datasets | pHV1 GC% | pHV1 size |
|---|---|---|---|---|---|---|---|---|
| **FS100** | *H. volcanii* | Δ*rnhA* Δ*rnhB* | 417 | 508 | DraI, EcoRV, SspI | 1 | 55.5 | 86 kb |
| **FS101** | *H. volcanii* | WT | 6 | 8 | | | | |
| **FS102** | *H. volcanii* | Δ*rnhA* Δ*rnhB* Δ*fen1* | 7 | 7 | | | | |
| **FS217** | *H. volcanii* | Δ*rnhA* | 197 | 262 | Fragmentase + MspA1I, HpyCH4V | 2 | | |
| **FS218** | *H. volcanii* | Δ*rnhB* | 33,266 | 169,298 | | | | |
| **FS219** | *H. volcanii* | Δ*rnhA* Δ*rnhB* | 2,611 | 3,079 | | | | |
| **FS223** | *H. volcanii* | Δ*rnhA* Δ*rnhB* Δ*fen1* | N/A | N/A | | | | |
| **FS220** | *H. volcanii* | WT | 81 | 107 | | | | |

| Library | Organism | Genotype | Genomic coordinates | Occurrences | Restriction Enzymes | ribose-seq datasets | pHV3 GC% | pHV3 size |
|---|---|---|---|---|---|---|---|---|
| **FS100** | *H. volcanii* | Δ*rnhA* Δ*rnhB* | 1,480 | 1,754 | DraI, EcoRV, SspI | 1 | 65.6 | 438 kb |
| **FS101** | *H. volcanii* | WT | 9 | 9 | | | | |
| **FS102** | *H. volcanii* | Δ*rnhA* Δ*rnhB* Δ*fen1* | 1,278 | 1,601 | | | | |
| **FS217** | *H. volcanii* | Δ*rnhA* | 958 | 1,225 | Fragmentase + MspA1I, HpyCH4V | 2 | | |
| **FS218** | *H. volcanii* | Δ*rnhB* | 202,905 | 1,003,649 | | | | |
| **FS219** | *H. volcanii* | Δ*rnhA* Δ*rnhB* | 17,158 | 20,906 | | | | |
| **FS223** | *H. volcanii* | Δ*rnhA* Δ*rnhB* Δ*fen1* | 2,264 | 2,929 | | | | |
| **FS220** | *H. volcanii* | WT | 506 | 603 | | | | |

| Library | Organism | Genotype | Genomic coordinates | Occurrences | Restriction Enzymes | ribose-seq datasets | pHV4 GC% | pHV4 size |
|---|---|---|---|---|---|---|---|---|
| **FS100** | *H. volcanii* | Δ*rnhA* Δ*rnhB* | 2,139 | 2,549 | DraI, EcoRV, SspI | 1 | 61.7 | 636 kb |
| **FS101** | *H. volcanii* | WT | 18 | 22 | | | | |
| **FS102** | *H. volcanii* | Δ*rnhA* Δ*rnhB* Δ*fen1* | 2,017 | 2,559 | | | | |
| **FS217** | *H. volcanii* | Δ*rnhA* | 955 | 1,219 | Fragmentase + MspA1I, HpyCH4V | 2 | | |
| **FS218** | *H. volcanii* | Δ*rnhB* | 219,219 | 848,154 | | | | |
| **FS219** | *H. volcanii* | Δ*rnhA* Δ*rnhB* | 18,874 | 22,526 | | | | |
| **FS223** | *H. volcanii* | Δ*rnhA* Δ*rnhB* Δ*fen1* | 2,669 | 3,557 | | | | |
| **FS220** | *H. volcanii* | WT | 588 | 751 | | | | |

**Table S3. Strand incorporation values of ribose-seq (R2 datatsets). (**N/A stands for data missing after the ribose-map process)

| | | Forward | Reverse |
|---|---|---|---|
| Hvo WT | Chr | 1,576 (48%) | 1,686(52%) |
| | pHV1 | 67 (63%) | 40(37%) |
| | pHV3 | 300 (50%) | 303 (50%) |
| | pHV4 | 416 (55%) | 335 (45%) |
| Hvo Δ*rnhA* | Chr | 2,832 (48%) | 3128 (52%) |
| | pHV1 | 150 (57%) | 112(43%) |
| | pHV3 | 620 (51%) | 605 (49%) |
| | pHV4 | 686 (56%) | 533 (44%) |
| Hvo Δ*rnhB* | Chr | 1,953,040 (49%) | 2,033,714 (51%) |
| | pHV1 | 95,743 (57%) | 73,555 (43%) |
| | pHV3 | 512,472 (51%) | 491,177 (49%) |
| | pHV4 | 483,679 (57%) | 364,475 (43%) |
| Hvo Δ*rnhA*Δ*rnhB* | Chr | 45,022 (49%) | 47,696 (51%) |
| | pHV1 | 1,742 (57%) | 1,337 (43%) |
| | pHV3 | 10,615 (51%) | 10,291 (49%) |
| | pHV4 | 11,891 (53%) | 10,635 (47%) |
| Hvo Δ*rnhA* Δ*rnhB* Δ*fen1* | Chr | 6,730 (47%) | 7,476(53%) |
| | pHV1 | N/A | N/A |
| | pHV3 | 1,381(47%) | 1,548 (53%) |
| | pHV4 | 2,082 (59%) | 1,475 (41%) |
| Tba WT | Chr | 27,271 (51%) | 26,594 (49%) |
| Tba Δ*rnhB* | Chr | 236,609 (49%) | 242,471 (51%) |

**Table S4. Strand incorporation values of ribose-seq (R1 datasets)**

| | | Forward | Reverse |
|---|---|---|---|
| Hvo WT | Chr | 26 (48%) | 28 (52%) |
| | pHV1 | 1 (12%) | 7 (88%) |
| | pHV3 | 3 (33%) | 6 (66%) |
| | pHV4 | 11 (50%) | 11 (50%) |
| Hvo Δ*rnhA* Δ*rnhB* | Chr | 3,034 (49%) | 3,171 (51%) |
| | pHV1 | 268 (52%) | 240 (48%) |
| | pHV3 | 956 (54%) | 798 (46%) |
| | pHV4 | 1,257 (49%) | 1,292 (51%) |
| Hvo Δ*rnhA* Δ*rnhB* Δ*fen1* | Chr | 2,702(50%) | 2,710 (50%) |
| | pHV1 | 6 (86%) | 1 (14%) |
| | pHV3 | 872 (54%) | 729 (46%) |
| | pHV4 | 1200 (47%) | 1359 (53%) |
| Tba WT | Chr | 638 (56%) | 503 (44%) |
| Tba Δ*rnhB* | Chr | 70,904 (52%) | 65,857 (48%) |

**Table S5. Rate of rNMP incorporation (%) for the R1 datasets.**

| Species | Genotypes | Genetic element | A | C | G | U |
|---|---|---|---|---|---|---|
| Hvo | WT | Chr | 23.7 | **31.2** | 13.6 | **31.6** |
| | | pHV1* | **64.0** | 10.2 | 0 | 25.6 |
| | | pHV3* | 0 | 25.4 | 25.4 | **49.3** |
| | | pHV4* | 16.5 | 23.9 | 10.2 | **49.4** |
| | Δ*rnhA* Δ*rnhB* | Chr | **38.9** | 26.7 | 29.3 | 5.1 |
| | | pHV1 | **39.0** | 27.7 | 26.1 | 7.1 |
| | | pHV3 | **39.3** | 26.7 | 27.9 | 6 |
| | | pHV4 | **38.4** | 28.8 | 26.9 | 5.8 |
| | Δ*rnhA* Δ*rnhB* Δ*fen1* | Chr | **40.6** | 26.7 | 25.8 | 6.8 |
| | | pHV1* | 16.1 | 42.9 | 14.3 | 32.3 |
| | | pHV3 | **42.7** | 38.8 | 12.9 | 6.6 |
| | | pHV4* | **42.1** | 27.9 | 22.2 | 7.8 |
| Tba | WT | Chr | 23.2 | **38.9** | 14.5 | 23.2 |
| | Δ*rnhB* | Chr | 29.1 | **45.5** | 24.6 | 0.9 |

*these values should be interpreted with caution because of the low global levels of ribonucleotide incorporation (see Table S2)

### Supplementary experimental procedures

#### Alignment of the amino acid sequences

Multiple sequence alignments of type 1 and 2 RNases H were computed by T-coffee (1) with four representative RNase H proteins from each kingdom of life whose sequence diversity were chosen to best illustrate the sequence variability. Structure-based sequence alignments were created by ESPript 3.0 (https://espript.ibcp.fr) (2).

#### Protein structure predictions and analyses

Protein structure predictions were created using nf-colabfold pipeline developed by SeBiMER, Ifremer's Bioinformatics Platform (https://sebimer.ifremer.fr/). nf-colabfold (https://gitlab.ifremer.fr/bioinfo/workflows/nf-colabfold) is a free and open-source Nextflow pipeline (3) capable of running ColabFold (4) on a mixed CPU/GPU cluster. MMseqs2 software (through colabfold_search tool) is executed on standard multi-core computing nodes, while AlphaFold software (though colabfold_batch tool) is executed on Nvidia GPU computing nodes. For the purpose of this study, release 1.3.0 of ColabFold has been used. All computations were undergone on the DATARMOR supercomputer hosted at Ifremer.

The predicted structures were superimposed and aligned using PyMol v2.4.1 software (Schrödinger) with two experimentally solved type 1 and 2 RNase H protein structures, which were representative of Eukaryotes and *Bacteria.* PyMol's *align* command was employed to visualize either the overlay of the overall protein structures or the superposition of the predicted active sites. The root-mean squared deviation (RMSD) were computed in PyMol, which performed 6 cycles of calculations on (X) aligned atoms (X pairs of C-alpha atoms) and, final RMSD (Å) values of atomic positions for (Y) atoms were reported (RMSD values (in Å) were given for Y atoms / X aligned atoms).

#### Supplementary references


1. Madeira, F., Pearce, M., Tivey, A.R.N., Basutkar, P., Lee, J., Edbali, O., Madhusoodanan, N., Kolesnikov, A. and Lopez, R. (2022) Search and sequence analysis tools services from EMBL-EBI in 2022. *Nucleic Acids Res*, **50**, W276-W279.
2. Robert, X. and Gouet, P. (2014) Deciphering key features in protein structures with the new ENDscript server. *Nucleic Acids Res*, **42**, W320-324.
3. Di Tommaso, P., Chatzou, M., Floden, E.W., Barja, P.P., Palumbo, E. and Notredame, C. (2017) Nextflow enables reproducible computational workflows. *Nat Biotechnol*, **35**, 316-319.
4. Mirdita, M., Schutze, K., Moriwaki, Y., Heo, L., Ovchinnikov, S. and Steinegger, M. (2022) ColabFold: making protein folding accessible to all. *Nat Methods*, **19**, 679-682.
